# PARK7/DJ-1 promotes pyruvate dehydrogenase activity and maintains Treg homeostasis

**DOI:** 10.1101/2019.12.20.884809

**Authors:** Egle Danileviciute, Ni Zeng, Christophe Capelle, Nicole Paczia, Mark A. Gillespie, Henry Kurniawan, Djalil Coowar, Daniela Maria Vogt Weisenhorn, Gemma Gomez Giro, Melanie Grusdat, Alexandre Baron, Coralie Guerin, Davide G. Franchina, Cathy Léonard, Olivia Domingues, Sylvie Delhalle, Wolfgang Wurst, Jens Christian Schwamborn, Rejko Krüger, Jeff Ranish, Dirk Brenner, Carole L. Linster, Rudi Balling, Markus Ollert, Feng Q. He

## Abstract

Pyruvate dehydrogenase (PDH) is the gatekeeper enzyme into the tricarboxylic acid (TCA) cycle. Here we show that PARK7/DJ-1, a key familial Parkinson’s disease (PD) gene, is a pacemaker controlling PDH activity in CD4 regulatory T cells (Tregs). DJ-1 bound to PDH-E1 beta (PDHB), inhibiting the phosphorylation of PDH-E1 alpha (PDHA), thus promoting PDH activity and oxidative phosphorylation (OXPHOS). *Dj-1* depletion impaired Treg proliferation and cellularity maintenance in older mice, increasing the severity during the remission phase of experimental autoimmune encephalomyelitis (EAE). The compromised proliferation and differentiation of Tregs in *Dj-1* knockout mice were caused via regulating PDH activity. These findings provide novel insight into the already complicated regulatory machinery of the PDH complex and demonstrate that the DJ-1-PDHB axis represents a potent target to maintain Treg homeostasis, which is dysregulated in many complex diseases.

## Introduction

Mitochondria are the organelle providing cellular bioenergy through tricarboxylic acid (TCA), oxidative phosphorylation (OXPHOS) and fatty acid oxidation. Compelling evidence shows that mitochondrial dysfunction contributes to several major neurological diseases such as Parkinson’s disease (PD) ^1,2^ and Alzheimer’s disease ^3^, but also to diseases of the gastrointestinal tract ^4^, diabetes ^5^ and cancer ^6^. In PD patients, a particular deficiency in mitochondrial complex I activity or general mitochondrial functions has been described ^7,8^. The activity and maintenance of the mitochondrial complex I is regulated by DJ-1/PARK7, a key risk gene for early-onset familial PD ^9-11^, via binding to its subunits ^12^. As a consequence, murine or human cells that are devoid of DJ-1/PARK7 expression showed mitochondrial defects affecting either mitochondrial respiration or integrity ^13-15^. Our current understanding of the functions of DJ-1 in mitochondria is essentially based on immortalized neurectodermal cancer cell lines and on primary or derived neurons ^16,17^. However, due to the existence of cell type- and tissue-specific protein interactions and differences in mitochondrial organelle number ^18^, these studies do not allow to extrapolate whether DJ-1 is also involved in the regulation of mitochondrial functions in non-neuronal cells such as in cells of the immune system, where mitochondrial dysfunction has been linked to the autoimmune diseases multiple sclerosis and rheumatoid arthritis ^19,20^. Recently, significant progress has already been made in understanding the role of *Dj-1* in innate immune cells, e.g., macrophages, where *Dj-1* interacts with p47phox, a subunit of the NADPH oxidase ^21,22^. As DJ-1 protects dopaminergic neurons from oxidative stress-induced cell death, the influence of DJ-1 on brain resident immune cells has also been investigated ^23^. Deletion of *Dj-1* indeed enhances activation of astrocytes and microglia, possibly through a STAT signaling pathway ^24^.

A potential contribution of the immune system to the pathogenesis of PD has been postulated for some time ^25,26^. As there is a complete penetrance of early-onset PD among individuals with homozygous DJ-1 loss-of-function mutations, we hypothesized that assessing the functions of DJ-1 in immune cells could provide valuable information on the impact of the molecular function of DJ-1 in cells that have a potential relevance for PD pathogenesis beyond neurons. So far, it still remains elusive whether and how DJ-1 is involved in the regulation of mitochondrial energetic metabolism on a molecular level and whether this could be potentially important for the physiological role of adaptive immune cells such as subsets of CD4^+^ T cells that have been associated with the pathogenesis of PD ^27^. While DJ-1 was already linked with the differentiation of inducible regulatory T cells (iTregs), no influence of DJ-1 on natural Tregs (nTregs) could be observed ^28^. Tregs are critical immune cells maintaining immune homeostasis and self-tolerance in both physiological and pathological conditions, such as in autoimmune diseases ^29-33^. Previous studies on DJ-1 function in Tregs were done by using young adult mice only^28^. Aging, however, is the most important risk factor for PD ^34-36^. Furthermore, a growing body of evidence showed a possible association between PD and autoimmunity ^37,38^. Thus, we hypothesized that the effects of DJ-1 on nTregs might become only evident during the aging process.

We undertook this study to examine the effects of DJ-1, a major genetic risk factor for early onset PD, on the CD4+ T cell function. Our data provide thorough evidence that DJ-1 is crucial in maintaining the cellularity of nTregs only in older adult mice, while no effect was seen in young adult mice. The mechanistic studies illustrate that DJ-1 interacts with pyruvate dehydrogenase (lipoamide) beta (PDHB), a subunit of the PDH super-complex, preferentially in CD4^+^CD25^+^Tregs, but not in CD4^+^CD25^-^ T cells. Furthermore, our results show that DJ-1 serves as a pacemaker governing the activity of PDH, the central metabolic enzyme linking the glycolytic pathway and the TCA cycle ^39,40^, preferentially in Tregs. Thus, by using both, *Dj-1* knockout (KO) mice and primary human T cells, we describe a previously unrecognized molecular mechanism through which DJ-1 critically regulates bioenergetic functions in mitochondria via the metabolic gatekeeper enzyme PDH. *Dj-1* depletion impaired Treg proliferation and maintenance of Treg cellularity in older mice, thereby increasing the severity of induced autoimmunity during the remission phase of experimental autoimmune encephalomyelitis (EAE). Thus, our data provide evidence that DJ-1 is an important component acting with PDH in Treg homeostasis, which contributes to an immune phenotype that becomes only apparent in adult mice at older age and that can be phenocopied through PDH inhibition in Tregs.

## Results

### Identification of *DJ-1* as a potential key gene in Tregs

Starting from our published dataset of high-time-resolution (HTR) time-series transcriptome data during the first 6 h of T cell receptor (TCR) stimulation ^41^, we found that *DJ-1* was the most highly expressed among 20 reported PD-related genes ^42^ in both unstimulated and stimulated human Tregs and Teffs (**Fig. 1a**). The immunoblot analysis from independent healthy donors confirmed the protein expression of *DJ-1* in both Tregs and Teffs (**Fig. 1b**, for Treg characterization refer to **Fig. EV1a-c**). Furthermore, by utilizing the so-called ‘queen-bee-surrounding principle’ network analysis strategy, derived from the ‘guilt-by-association’ concept ^43-45^, we analyzed the first-degree neighborhoods in the *DJ-1* subnetwork. However, the first-degree neighbors of *DJ-1* were not significantly enriched in a collection of 400 genes potentially related to T-cell function ^41^. Since the functions of the second-degree neighbors in the network might also be indicative of the roles of the given node ^43,44^, we further extended to scrutinize the subnetwork composed of second-order neighbors of *DJ-1*. Interestingly, *DJ-1* was then highly connected with well-known key players in Treg functions (**Fig. 1c**, *P=4.3E-12*, cumulative binomial distribution), such as *FOXP3, CTLA4, ICOS, GATA3*, and *CD44* ^30^. None of the other known PD genes were highly expressed in Tregs and Teffs and significantly surrounded by potential T-cell-related genes in the Treg-specific network (**Table EV1**). Together, this network analysis results predict that DJ-1 might play a role in Tregs that warrants further investigation of its potential impact on Treg function and cellularity.

**Figure 1.**
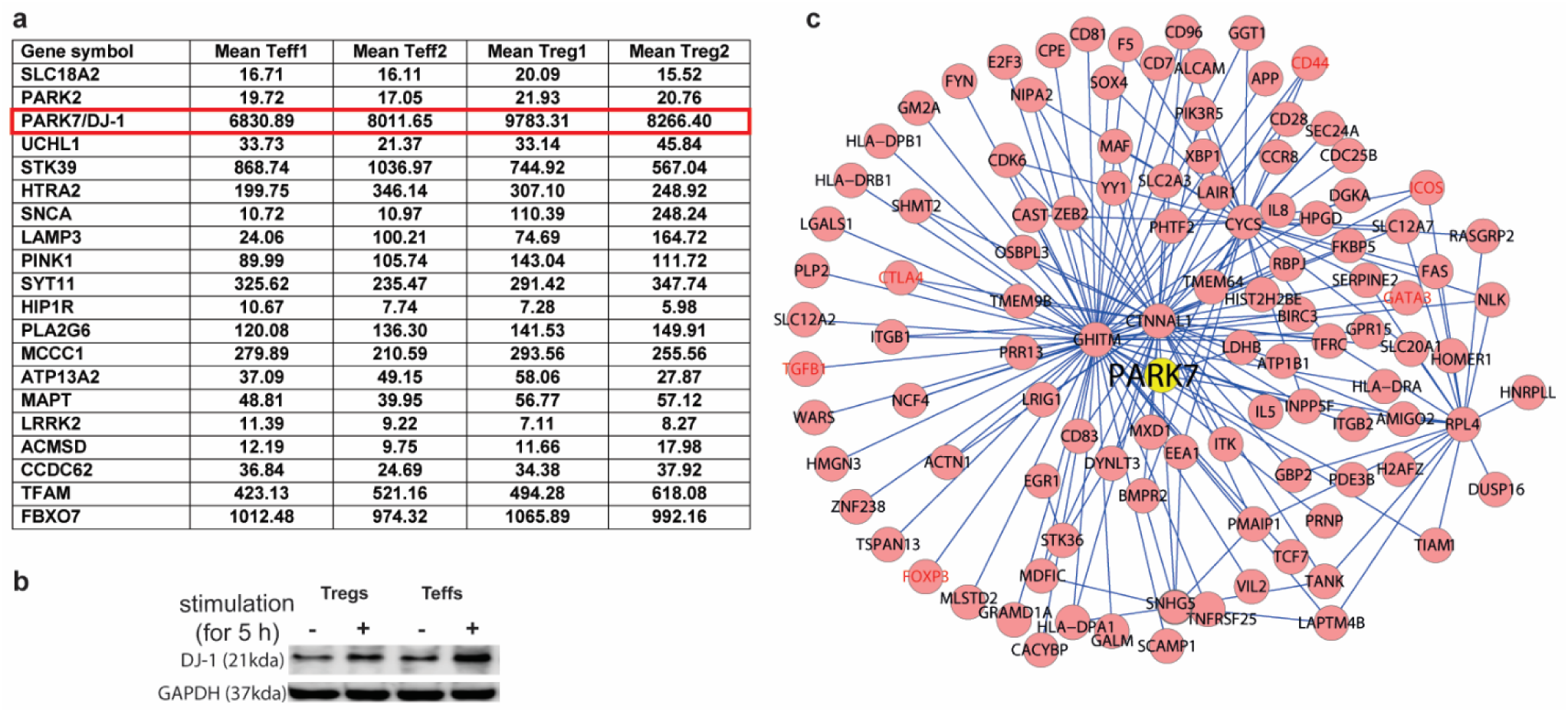
DJ-1 is highly expressed in human Tregs and connected with well-known Treg genes in Treg specific-correlation network. **a**, mean transcription expression of the known PD genes measured in the first 6 h of anti-CD3-/-CD28/IL2 stimulation assessed by our published high-time-resolution (HTR) time-series transcriptome (with 19 time points) ^41^. Treg1 and Treg2 (or Teff1 and Teff2) represent two independently repeated HTR time-series experiments for Tregs (or Teffs). DJ-1 is highlighted by red rectangle. **b**, DJ-1 protein expression in human primary Tregs and Teffs with or without anti-CD3/-CD28/IL-2 stimulation assessed by Western Blotting. **c**, The PARK7/DJ-1 subnetwork extracted from the constructed human Treg-specific correlation network. Each oval represents one gene. A selection of known key Treg genes were highlighted in red. Each line between DJ-1 and the other genes represents a correlation-based functional linkage and the other genes represents a correlation-based functional linkage.

### Dj-1 depletion impairs Treg cellularity maintenance in aged mice

To test whether DJ-1 maintains Treg cellularity as suggested above, we analyzed the Treg compartment in *Dj-1*^*-/-*^knockout (KO) mice as developed elsewhere ^46^. Since DJ-1 is one of the key PD familial genes and aging is the major risk factor of PD ^35,36^, we investigated both young adult and older mice. For both young adult mice (short as ‘young mice’, 8–12 wks) and older adult mice (short as ‘old mice’, ∼45 wks), the frequency of total CD4^+^ T cells in spleen and peripheral lymph nodes was not significantly changed between *Dj-1 KO* mice and *Dj-1*^+/+^ littermate control mice (WT, **Fig. 2a**). The frequency of total Tregs (CD4^+^FOXP3^+^ T cells, Tregs) among total CD4^+^ T cells was also not different between young *Dj-1 KO* mice and WT mice. Consistent with other reports ^47^, the frequency of Tregs among total CD4^+^ cells was increased with age in WT mice (**Fig. 2b**). Interestingly, the frequency of Tregs was significantly decreased in spleen and peripheral lymph nodes of old *Dj-1 KO* mice compared with that of age- and gender-matched WT mice (**Fig. 2b, c**). The percentages of CD4^+^CD25^+^ T cells among total CD4^+^ T cells showed similar changes as that of CD4^+^FOXP3^+^ T cells (**Fig. 2d, e**). The total absolute number of splenic Tregs was also significantly reduced in old *Dj-1 KO* mice compared with WT mice (**Fig. 2f**). The ratio between CD4^+^FOXP3^-^ Tconv and Tregs was increased in old but not young *Dj-1 KO* mice (**Fig. 2g**). As Helios is a marker distinguishing natural Tregs (nTregs) from induced Tregs (iTregs), we also analyzed expression of Helios. Interestingly, the percentages of Helios positive cells among CD4^+^FOXP3^+^ T cells also declined in old but not young *Dj-1* KO mice vs. WT mice (**Fig. 2h, i**). Consistent with reduced Tregs, the frequency of splenic IL-10-producing CD4^+^ cells following *in vitro* PMA/ionomycin stimulation was significantly decreased in old *DJ-1* KO mice vs. WT littermates (**Fig. 2j**). Furthermore, the age-dependent effect of the *Dj-1* loss on Treg cellularity was not simply caused by the potential differential expression of *Dj-1* between young and old WT mice (**Fig. 2k**). Despite the reduced Treg cellularity, old *Dj-1* KO mice up to the tested age did not develop spontaneous autoimmune phenotypes, possibly due to the compensation by imbalanced Tconv cell subsets (unpublished own data).

**Figure 2.**
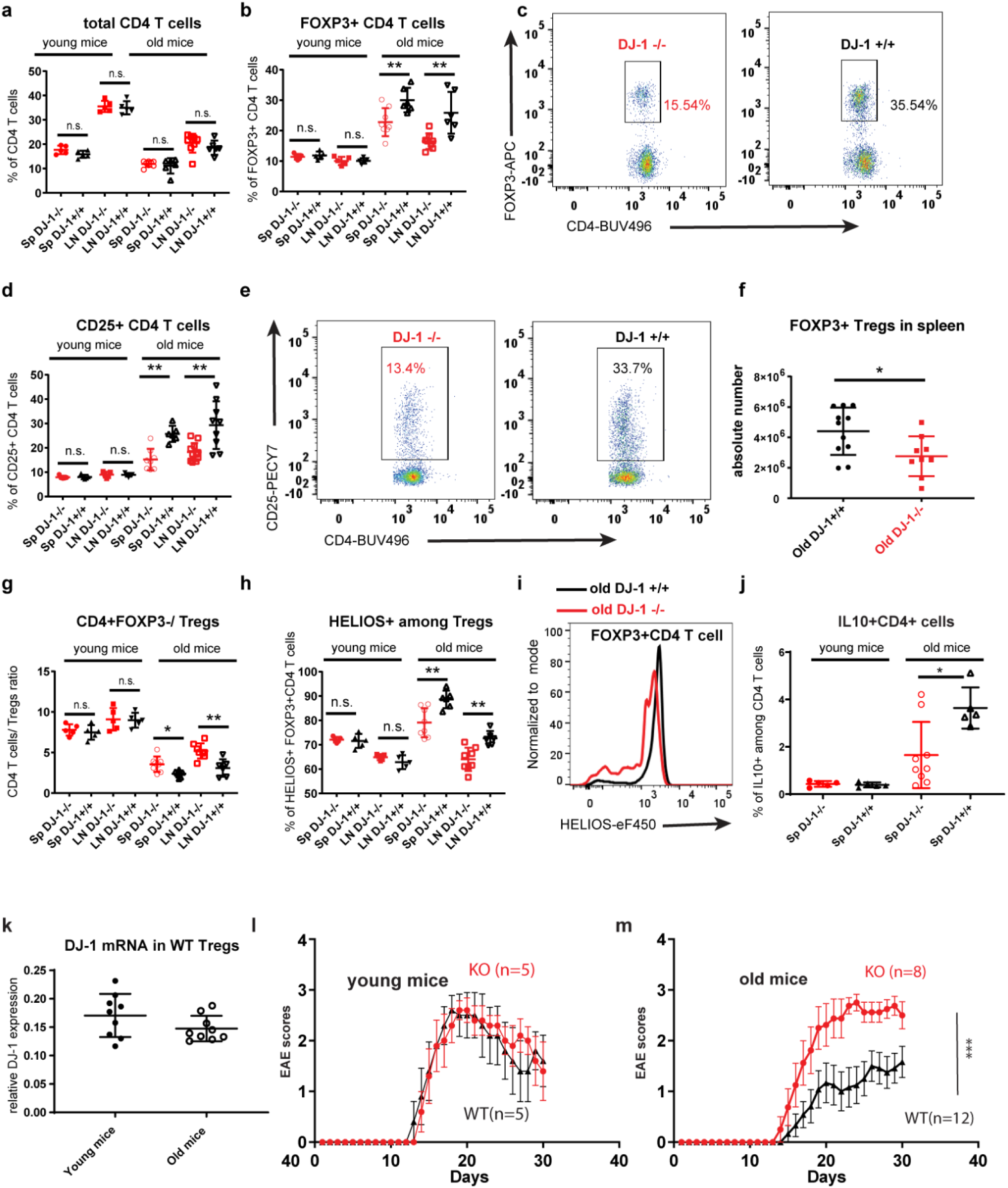
*Dj-1* depletion reduces Treg accumulation in old mice. **a**, Percentages of CD4+ T cells from spleen (Sp) or peripheral lymph nodes (LN) (young KO, n=5; young WT, n=5; old KO, n=8; old WT, n=6; for old mice, data pooled from 2 independent experiments). Each symbol represents one mouse unless otherwise mentioned. **b**, Percentages of FOXP3+ subsets among CD4 T cells. **c**, Representative flow-cytometry plots of FOXP3+ cells among total CD4 T cells in spleen of old WT or DJ-1 KO mice. **d**, Percentages of CD25+ subsets among CD4 T cells (young mice number as in **a**, but for old mice: spleen WT, n=6;spleen KO, n=8;LN WT, n=9; LN KO, n=11, data pooled from 2 independent experiments). **e**, Representative flow-cytometry plots of CD25+ cells among total CD4 T cells in spleen of old WT or DJ-1 KO mice. **f**, Number of FOXP3^+^CD4^+^ T cells in spleen of indicated genotype mice (estimated from total splenocytes, WT, n=9; KO, n=11, pooled data). **g**, The relative ratios between non-Treg cells and FOXP3^+^CD4^+^ Tregs. **h**, Percentages of Helios+ subsets among FOXP3^+^CD4^+^ Tregs. **i**, Representative histogram overlay of Helios expression among total FOXP3^+^CD4^+^ Tregs between old WT or DJ-1 KO mice. **j**, Expression of IL-10 in CD4 T cells after in-vitro stimulation using PMA/ionomycin for 5 h. **k**, qPCR quantification of Dj-1 expression in sorted Tregs from young or old WT mice (n=3/group and 3 technical replicates/mouse). Results represent at least four (**a-j**) and two (**k**,**l**,**m**) independent experiments. Data are mean± s.d. The P-values are determined by a two-tailed Student’s t-test. ns or unlabelled, not significant, *P<=0.05, **P<=0.01 and ***P<=0.001. **l, m**, EAE clinical scores on the indicated days following immunization with MOG35-55 in young (**l**) or old mice (**m**). The number of age- and gender-matched mice per group are indicated in the subfigures. Data are representative for young mice and a summary of 2 independent experiments for old mice groups. Young mice (∼12 wks, left) and old mice (∼45 wks, right).

Functional Tregs are critical for the active phases (e.g., remission) but not the onset of experimental autoimmune encephalomyelitis (EAE) ^48^, a murine model of the human autoimmune disease multiple sclerosis. We therefore evaluated the *in vivo* effects of reduced Treg accumulation in *Dj-1 KO* mice by using the myelin oligodendrocyte glycoprotein (MOG35-55)-induced EAE model ^49^. In agreement with not seeing a significant difference in the Treg frequency between young *Dj-1 KO* mice and their WT littermates, no obvious difference in the clinical scores of different disease phases was observed (**Fig. 2l**). Interestingly, old *Dj-1 KO* mice relative to their age- and gender-matched WT controls exhibited deteriorated EAE symptoms during the Treg-dependent disease remission phase (**Fig. 2m**). These data suggested an impairment of total *in vivo* immunosuppression capability in *Dj-1 KO* mice attributable to a decreased number of Tregs.

The decreased Treg accumulation in aged mice could be caused by an impairment either in Treg development or Treg homeostatic proliferation. To test these two possibilities, we first assessed the thymic Treg compartment in 4-wk-old mice as thymocytes peak in mice aged at 4-6 weeks and gradually evolve afterwards ^50^. In 4-wk-old mice, we did not observe any difference between *Dj-1 KO* mice and WT mice in total thymocytes, single-positive CD4 T cells (CD4 SP), single-positive CD8 T cells (CD8 SP) or the percentages of total FOXP3+ Tregs among CD4 SP (**Fig. 3a-d**). We also did not find any difference in the four subsets determined by the combination of positive or negative expression of CD25 and FOXP3 (**Fig. 3e**). Neither Treg markers, nor proliferation or functional markers, such as Helios, Ki67 or PD-1, CTLA4 and GITR exhibited any difference between 4-wk-old *Dj-1 KO* and WT mice (**Fig. 3f-h**). These data suggest that *Dj-1* depletion does not influence Treg development in juvenile mice before thymic involution. As decreased Treg accumulation only appeared in aged *Dj-1 KO* mice, we also analyzed the cellular composition of thymocytes in old mice (∼45 wks). As thymic atrophy already started in 45-wk-old mice, the total thymocytes were dramatically reduced in old WT mice relative to 4-wk-old mice (**Fig. 3a**). However, the number of total thymocytes was not significantly affected when comparing old *Dj-1 KO* mice with WT mice at the tested age (**Fig. 3a**). No significant difference was observed between old KO and WT mice with respect to the frequency of CD4 SP, CD8 SP and the frequency of total Tregs among CD4 SP (**Fig. 3b-d**). Interestingly, *Dj-1* ablation selectively decreased the frequency of Helios-expressing cells among FOXP3^+^CD4^+^ T cells (**Fig. 3g**) and modestly but significantly reduced homeostatic proliferation of FOXP3^+^CD4^+^ T cells (**Fig. 3h, i**). Therefore, *Dj-1* depletion did not affect Treg development, neither in 4-wk-old nor in aged mice, indicating that the observations in peripheral lymphoid tissues of aged mice might be caused by deficiency in Treg homeostatic proliferation.

**Figure 3.**
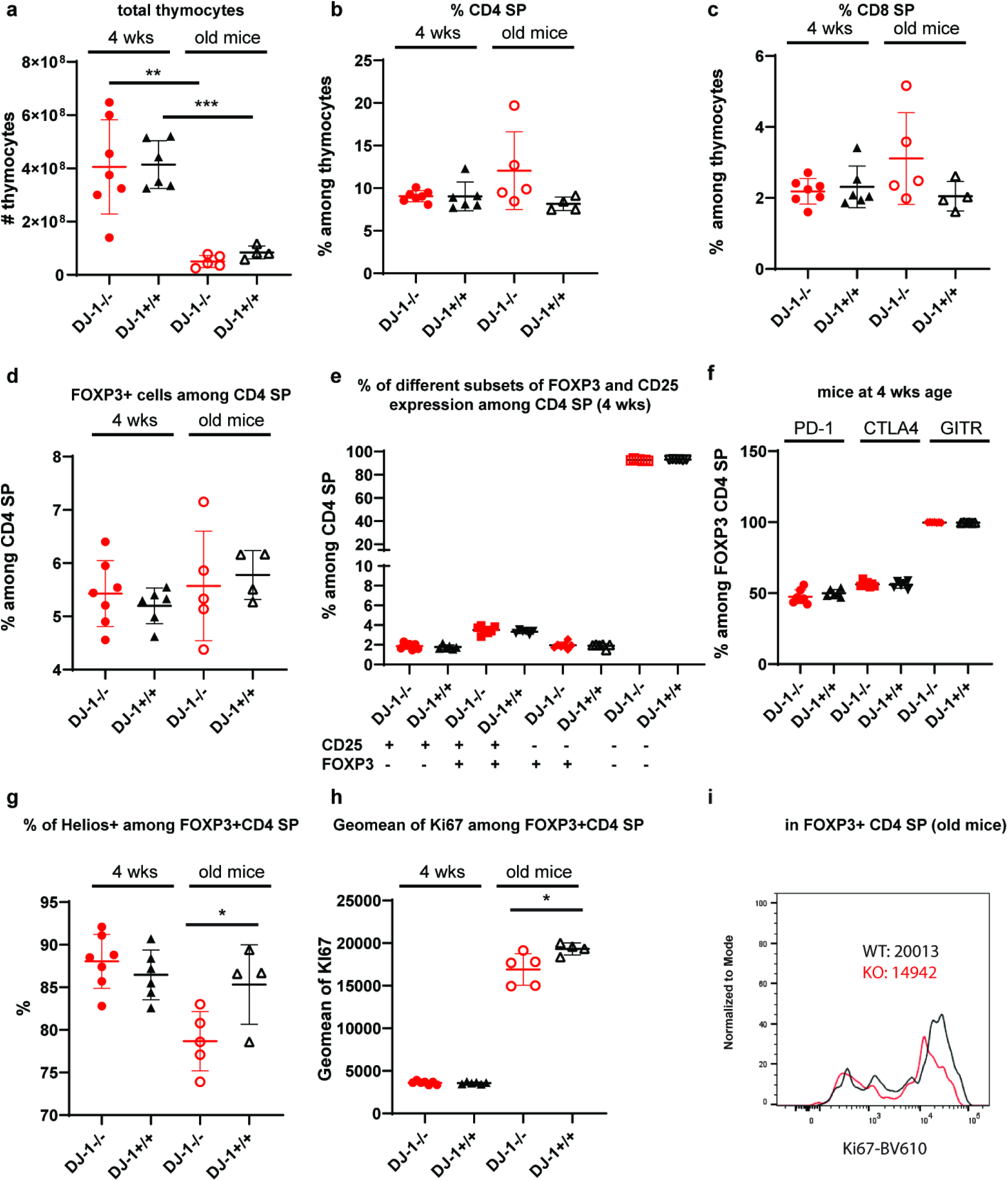
*Dj-1* ablation does not affect thymic Treg development independent of age. **a**, Total thymocytes of 4-wk-old or old *Dj-1 KO* and WT mice (young KO, n=7; young WT, n=6;old KO, n=5;old WT, n=4). **b, c**, Percentages of CD4 single positive (SP) cells (**b**) and CD8 SP (**c**) among thymocytes. **d**, Percentages of FOXP3^+^ among CD4SP cells. **e**, Percentages of PD-1, CTLA4 and GITR among FOXP3^+^CD4SP cells in 4-wk-old mice **f**, Percentages of the four subsets defined by the presence or absence of FOXP3 and CD25 among CD4SP cells in 4-wk-old mice. **g**, Percentages of Helios^+^ among FOXP3^+^CD4SP cells. **h**, Geometric mean of Ki67 among FOXP3^+^CD4SP cells. **i**, Representative histogram overlay of Ki67 expression among total FOXP3^+^CD4+SP cells between old WT or *Dj-1 KO* mice. Results represent three (**a-i**) independent experiments. Data are mean± s.d. The P-values are determined by a two-tailed Student’s *t*-test. ns or unlabeled, not significant, *P<=0.05, **P<=0.01 and ***P<=0.001.

### Dj-1 depletion impairs Treg proliferation and activation in aged mice, but not Treg suppessor function

The finding of a decreased frequency of Tregs in old *Dj-1 KO* mice prompted us to seek for the underlying molecular pathways mediated by DJ-1. We performed transcriptomic analysis of sorted *Dj-1 KO* and WT Tregs from spleen of old mice. As expected, the upregulated Treg-signature genes were expressed much higher than Tconv-signature genes ^51^, indicating a highly-purified cell sorting (**Fig. 4a**). Consistent with the view of DJ-1 as a transcriptional coactivator ^52^, *Dj-1* deficient Tregs exhibited much larger fractions of downregulated genes compared with those of upregulated genes (**Fig. 4b**). In old *Dj-1 KO* Tregs, in line with reduced Ki67 staining (**Fig. 3h, i**), the genes involved in cell cycle and proliferation were significantly enriched among the downregulated list (**Fig. 4c, d**). Moreover, the loss of *Dj-1* in Tregs of old mice reduced the transcription of many key Treg genes, such as *Tnfrsf1b, Tnfrsf8* and *Itgae* ^51^, as well as many of the MHC II-related genes, which are involved in the early-contact-dependent suppression ^53^ (**Fig. 4c**). Many of these MHC II-related genes were also observed in the human Treg-specific correlation network surrounding *DJ-1* (**Fig. 1c**). The MHC II genes are reported to be expressed in activated T cells of many species, but are not expressed in activated murine T cells ^54^, possibly because this aspect has never been analyzed in depth in aged mice. The observation of decreased MHC II genes is in agreement with the decreased homeostatic expression of activation markers in Tregs (**Fig. 4e, f**), such as GITR and ICOS, in old *Dj-1 KO* mice vs. WT mice. This unbiased transcriptome analysis demonstrates that *Dj-1* deficiency leads to a downregulation of pathways mediating Treg homeostatic proliferation and activation, which accumulatively contributes to the reduced Treg compartment observed in old *Dj-1* KO mice. In conclusion, DJ-1 is critical for maintaining Treg cellularity via regulating their activation and proliferation, which becomes evident only in aged mice.

**Figure 4.**
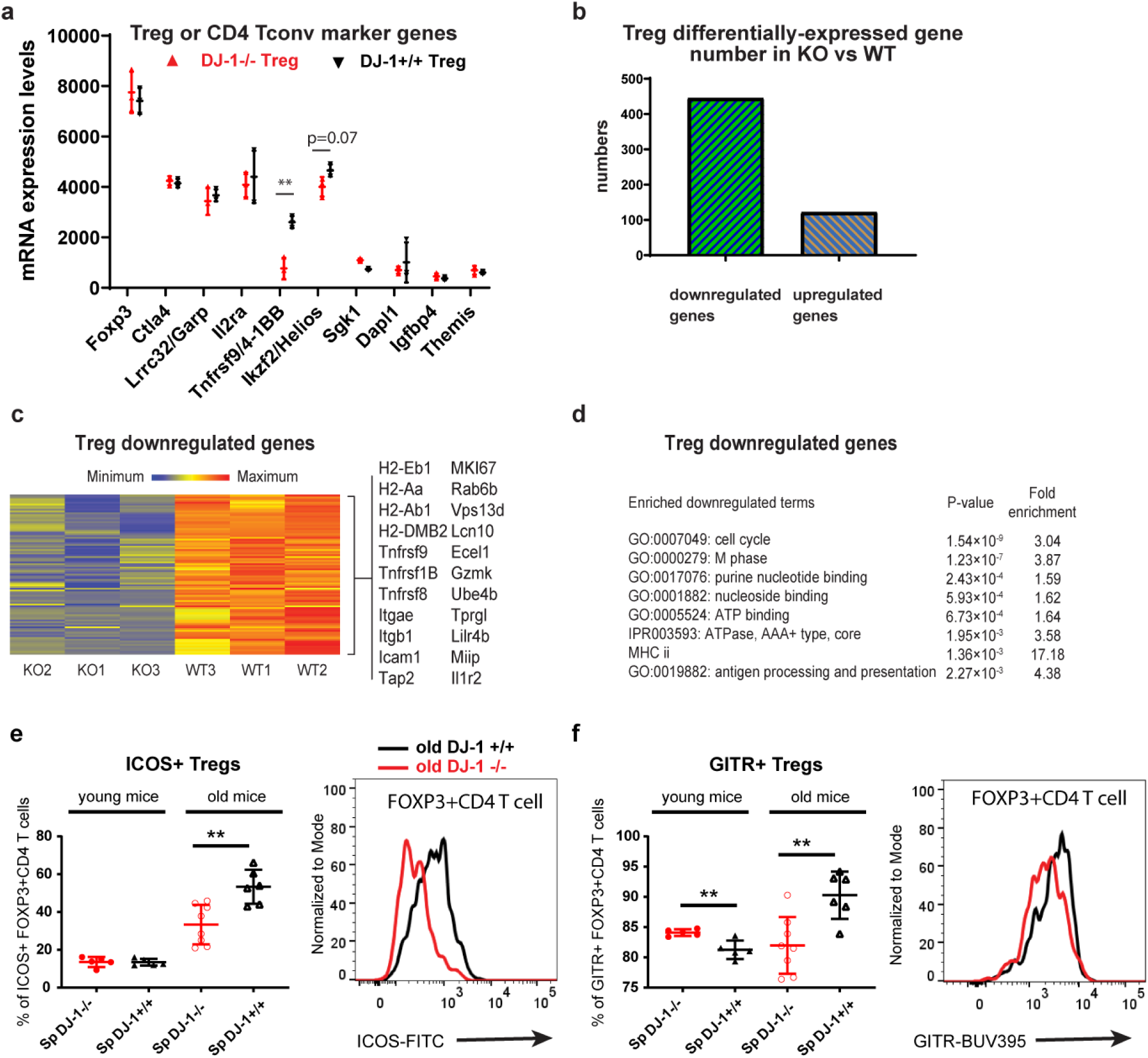
*Dj-1* depletion impairs Treg proliferation and activation pathways in old mice. **a**, mRNA expression of upregulated Treg-signature and Tconv-signature genes in Tregs freshly isolated from old *Dj-1 KO* and WT littermates. **b**, The number of differentially-expressed up- and down-regulated genes in Tregs. **c**, Heatmap showing relative abundance of selected differentially downregulated genes in Tregs from old *Dj-1* KO mice vs. the age-matched WT littermates. n=3 mice/group **d**, The selected significantly enriched GO terms among downregulated genes in Tregs from old *Dj-1* KO mice vs. the age-matched WT littermates. **e, f**, Percentages of ICOS+ (**e**) or GITR+ (**f**) subsets among FOXP3+CD4+ T cells (left) and representative histogram overlay of ICOS expression among total FOXP3+CD4+ Tregs between old WT or *Dj-1 KO* mice (right) (young KO, n=5; young WT, n=5;old KO, n=8;old WT, n=6). Results are the summary of the microarray datasets. Data are mean± s.d. The P-values are determined by a two-tailed Student’s *t*-test. ns or unlabelled, not significant, *P<=0.05, **P<=0.01 and ***P<=0.001.

As outlined earlier by the computational correlation network analysis, we also tested whether *Dj-1* plays a critical role in controlling Treg suppressor function (TSF). When we analyzed TSF from young mice *in vitro* using Tregs, CD4 T conventional cells (Tconv) and feeder cells in co-culture suppressive assays, no significant difference was observed between young *Dj-1 KO* and WT mice (**Fig. EV2a**), consistent with a previous study ^28^ in which the authors also investigated the suppressor function of DJ-1-KO Treg, but only focused on young mice. Interestingly in old *Dj-1 KO* mice relative to WT mice, *in vitro* TSF was also not compromised (e.g., at the ratio of 1:1, **Fig. EV2b**), but even slightly augmented with a higher ratio between Tconv and Tregs (e.g., at the ratio of 4:1, **Fig. EV2b**). To further characterize the Tregs, we investigated the protein expression of known key Treg effector molecules, such as FOXP3, CTLA4, and IL2RA (CD25) in Tregs of old DJ-1 mice. Although the fraction of CTLA4 positive cells among old *Dj-1 KO* Tregs was not different compared with that among WT Tregs (**Fig. EV2c**), the expression levels of CTLA4 among CTLA4 positive cells were modestly, but significantly increased in spleen (**Fig. EV2c**). Furthermore, FOXP3 and CD25 expression levels were also modestly increased in spleen of old *Dj-1*-KO individual Tregs (**Fig. EV2d, e**). Therefore, our data demonstrate that DJ-1 is critical for maintaining Treg cellularity, but dispensable for Treg suppressor function.

### *DJ-1* promotes PDH activity preferentially in Tregs

To further identify the detailed underlying molecular mechanisms through which DJ-1 maintains Treg cellularity, we next examined the binding partners of DJ-1 in resting and activated human primary nTregs by coimmunoprecipitation (IP)-mass spectrometry (MS) analysis. We identified around 20 novel potential binding partners of DJ-1 in human nTregs by employing a combination of five strict filtering criteria (**Fig. 5a, Table EV2**, refer to Methods). Interestingly, five proteins regulating RNA splicing, namely, HNRNPH3, PRPF8, KHSRP, SKIV2L2 and SART1, bound to DJ-1 preferentially in the stimulated Tregs and Teffs compared with those in the resting cells (P=2.8E-4, **Fig. 5a, Table EV2**). This is consistent with the dynamic binding behavior of DJ-1 to RNAs ^11^, which binds to RNAs in a resting state and then dissociates following activation-driven oxidative stress as shown in a neuroblastoma cell line. Our results also confirmed 11 known DJ-1 binding partners identified previously from other cell types (**Table EV3**, refer to Methods), demonstrating the reliability of our technique.

**Figure 5.**
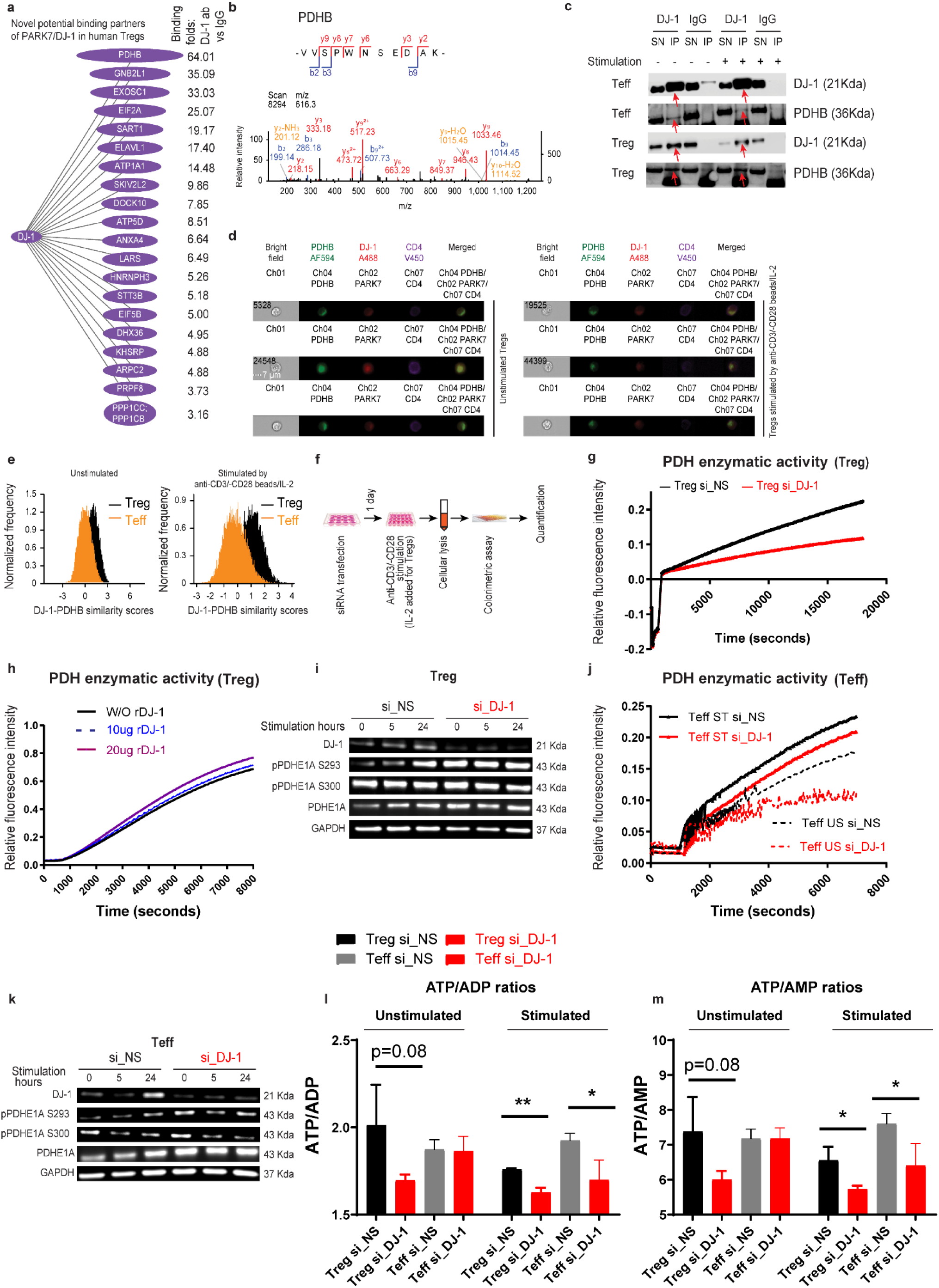
DJ-1 promotes PDH activity selectively in Tregs. **a**, Potential novel binding partners of DJ-1 as identified by the co-IP-MS analysis. The binding enrichment of the corresponding protein to anti-DJ-1 ab relative to IgG control in human primary Tregs is presented on the plot. The width of the oval is relatively correlated to the binding specificity. **b**, Representative MS/MS spectrum for a peptide used to identify PDHB. Observed b- and y-fragment ions are labeled within the spectrum, and summarized in the peptide sequence shown above. **c**, Validation of DJ-1 bound to PDHB preferentially in human Tregs relative to Teffs by Co-IP-immunoblotting (Co-IP-IB) analysis. The red arrow indicates DJ-1 or PDHB band from anti-DJ-1 IP elution. Co-IPs were performed with anti-DJ-1 or IgG in the lysates of primary human Tregs/Teffs with or without anti-CD3/-CD28/IL-2 stimulation for 5 hours. To independently validate the cobinding discovery from the MS analysis, the cells used in the immunoprecipitation-immunoblotting and colocalization analysis were isolated from different healthy donors. SN, supernatant; IP, eluted complexes pulled out by anti-DJ-1 or IgG control ab. **d**, Representative single cell images using Imagestream. While scale bar, 7um; original magnification x60. **e**, DJ1/PARK7 and PDHB are preferentially co-localized in Tregs relative to Teffs. The similarity scores are trans-location similarity scores showing to which degree DJ-1/PARK7 is translocated to the specific area, where PDHB is located. **f**, Schematic on how the enzymatic assay of PDH was performed. **g**, Representative data of PDH enzymatic activity of stimulated human Tregs treated with specific siRNA against DJ-1 or non-specific scambled siRNA. **h**, PDH enzymatic activities of stimulated human Tregs without (W/O) or with various amount of addition of recombinant human DJ-1 protein. **i, k**, Site-specific phosphorylation and expression of PDHA in Tregs (**i**) or Teffs (**k**) transfected with or without DJ-1 knockdown followed by different stimulation periods. **j**, PDH enzymatic activities of unstimulated (US) or stimulated (ST) Teffs with or without *DJ-1* knockdown. **l, m**, Ratios between intracellular ATP and ADP (**l**) or between intracellular ATP and AMP (**m**) in Tregs or Teffs with or without *DJ-1* knockdown, and with or without stimulation. Results represent four (**c, g**), three (**d, e, i, j, k, l, m**) and one (**a, b**) independent experiments. Data are mean± s.d. The P-values are determined by a two-tailed Student’s *t*-test. ns or unlabelled, not significant, *P<=0.05, **P<=0.01 and ***P<=0.001.

To prioritize and test the most promising binding candidates, we ranked the top 20 novel binding partners based on their preferential binding to DJ-1 relative to the IgG control (**Fig. 5a**). Unexpectedly, the top-ranked candidate was PDHB (pyruvate dehydrogenase [lipoamide] beta). Furthermore, PDHB also occupied the highest fold change in binding affinity between unstimulated and stimulated Treg samples in the MS analysis (**Table EV2**). Following the DJ-1 Co-IP-MS identification of PDHB (**Fig. 5a, b**), we confirmed that DJ-1 bound to PDHB by coIP-immunoblotting analysis and observed that the DJ-1 binding was more predominant in Tregs than in Teffs (**Fig. 5c**). Moreover, using ImageStream analysis, we quantitatively demonstrated that DJ-1 preferentially colocalized with PDHB in Tregs rather than in Teffs, with or without anti-CD3/-CD28/IL-2 stimulatory beads or antibodies (**Fig. 5d, e**). Coincidently, an available murine microarray database, analyzing the mRNA coexpression in different integrated datasets in a cell-type-specific manner, demonstrated that *Dj-1* and *Pdhb* are significantly correlated in murine Tregs (**Fig. EV3a**), but not in other CD4 T-cell subsets, such as effector memory CD4 T (CD4 Tem) cells, Th1 or Th2 cells ^55^ (**Fig. EV3b**). This finding is not only in line with the reported role of Dj-1 in glucose homeostasis of pancreatic islets ^56^, but also suggests a novel molecular pathway that connects Treg function or maintenance with metabolism. Taken together, our data from cobinding, colocation and coexpression strongly support that DJ-1 binds to PDHB preferentially in Tregs relative to Teffs.

To further investigate whether the physical binding between DJ-1 and PDHB might result in any functional consequences, we checked PDH enzymatic activity. Interestingly, knocking down *DJ-1* significantly reduced PDH enzymatic activity in human Tregs (**Fig. 5g**), while adding recombinant human DJ-1 protein to the T-cell lysates promoted PDH activity in a high-dose-dependent manner (**Fig. 5h**). The requirement for a high-dose of recombinant DJ-1 protein to further enhance PDH activity is most likely due to the high endogenous expression of DJ-1. Since the PDH enzyme is mainly activated through site-specific dephosphorylation of three sites (Ser232, Ser293 and Ser300) on the E1 alpha subunit (PDH-E1A, also known as PDHA) ^40^, we measured the site-specific phosphorylation in human Tregs. Consistent with the impact on PDH enzymatic activity, knocking down *DJ-1* in Tregs significantly increased the phosphorylation levels of Ser293 but not another tested site (Ser300) (**Fig. 5i**). In line with the weaker but detectable binding between DJ-1 and PDHB in Teffs relative to Tregs (**Fig. 6c**), knocking down *DJ-1* in Teffs modestly dampened PDH activity (**Fig. 5j**) and enhanced Ser293 phosphorylation on PDHA (**Fig. 5k**). Notably, a stronger effect of DJ-1 on the phosphorylation of Ser293 was observed in the unstimulated Tregs and Teffs (**Fig. 5i, k**) compared to the stimulated counterparts. This finding is in line with the preferential binding in unstimulated vs. stimulated T cells as determined by MS analysis (**Table EV2**). A stronger effect on PDH activity in resting T cells is also in agreement with the fact that following stimulation, T cells switch their energetic model from mainly OXPHOS to glycolysis ^57^, a situation in which T cells rely less on PDH activity. Taken together, these findings support the notion that binding of DJ-1 to PDHB promotes PDH enzymatic activity by inhibiting Ser293 phosphorylation levels of PDHA in Tregs.

**Figure 6.**
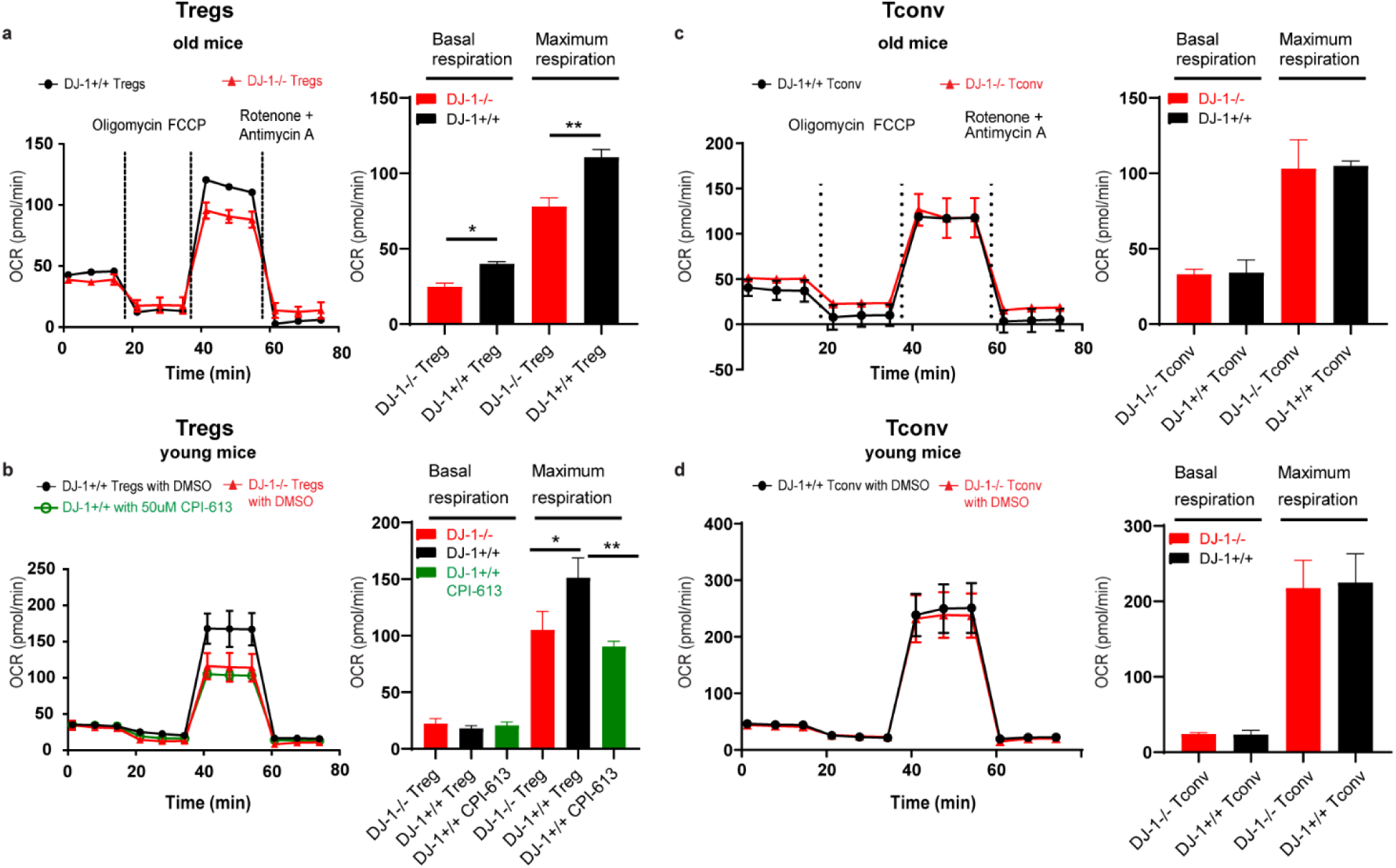
*Dj-1* depletion impairs OXPHOS selectively in Tregs. **a, b, c, d**, Oxygen consumption rate (OCR) of freshly isolated Tregs (**a**, old mice; **b**, young mice) and CD4 conventional cells (Tconv) (**c**, old mice; **d**, young mice) from *Dj-1 KO* and WT mice. left of each panel, representation plots following Mito stress test. right of each panel, quantitation of basal and maximum respiration under Mito stress test conditions. Results represent three (**a-d**) independent experiments. Data are mean± s.d. The P-values are determined by a two-tailed Student’s *t*-test. ns or unlabelled, not significant, *P<=0.05, **P<=0.01 and ***P<=0.001.

Reduced PDH activity following DJ-1 knockdown in Tregs could be also directly regulated by the expression levels of PDH kinases (*Pdks*) and phosphatases (*Pdps*) ^58^ as DJ-1 is also known as a transcription co-activator. We therefore analyzed those genes using the Treg transcriptomic data sorted from old *Dj-1 KO* and WT mice, but none of the *Pdp* genes showed significant difference at the transcriptional level (**Fig. EV4a**). Interestingly, the kinase gene *Pdk1* was upregulated while the other two kinase genes *Pdk2* and *Pdk3* were downregulated in old *Dj-1 KO* mice, three of which might compensate each other and contribute to the overall PDH activity in Tregs (**Fig. EV4a**). The change in PDH activity was not simply rendered by a change in mRNA of the genes encoding the PDH supercomplex ^59^ as no change was observed for any of these genes (**Fig. EV4b**). Thus, DJ-1 regulates the PDH activity in Tregs, not only via a physical interaction with PDHB, but also possibly in parallel via transcriptional regulation of PDH kinase genes. Nevertheless, to distinguish the contribution of each mechanism to PDH activity requires further investigation.

Since PDH is the critical TCA-cycle-entry enzyme and crucial for ATP generation, a partial knockdown of *DJ-1* in Tregs indeed decreased both the ATP/ADP and ATP/AMP ratios, with or without TCR stimulation (**Fig. 5l, m**). For stimulated Teffs, DJ-1 downregulation reduced both ratios (**Fig. 5l, m**). Therefore, these data demonstrate that DJ-1 also regulates ATP production in Tregs.

Having identified DJ-1 as a binding partner of PDHB, which consequently affects PDH activity, the critical gatekeeper enzyme entering the TCA cycle, we sought to further evaluate whether DJ-1 has any effect on OXPHOS of Tregs, as Tregs critically depend on OXPHOS ^60^. Particularly, we measured the mitochondrial reserve or spare respiratory capability (SRC) and maximum respiration. These measurements are used as a proxy for the extra mitochondrial capacity available to produce energy through OXPHOS under activation or cellular stress conditions, which is important for the long-term survival of the cell ^61^. Due to the loss of *Dj-1* and its binding with PDHB, an impaired SRC was indeed observed in Tregs of old *DJ-1 KO* mice (**Fig. 6a**), resulting in a lower bioenergetic production and consequently reduced proliferation (**Fig. 3h, i**). Interestingly, the baseline OXPHOS was only reduced in *Dj-1 KO* Tregs from old but not young mice (**Fig. 6a, b**), which might subsequently impair Treg homeostatic proliferation (**Fig. 3h, i**). However, the maximum respiration was already decreased in young *Dj-1 KO* Tregs (**Fig. 6b**), indicating that under stressful cellular or activation conditions, Treg survival or proliferation might be affected. The aging progression itself could be considered as a chronic inflammatory/activation process or model and could be the reason why we observed an impaired accumulation of Treg cellularity in aged, but not young *Dj-1 KO* mice. We did not observe any significant difference in the baseline OXPHOS and SRC of Tconv, neither in young nor old *Dj-1 KO* and WT mice (**Fig. 6c, d**). This is not only consistent with the current notion that Tregs rely on OXPHOS much more than do Tconv cells ^60,62^, but also is reflected by our observation that Tconv cellularity was not affected in both young and old *Dj-1 KO* mice (**Fig. 2a**). In short, DJ-1 binds to PDHB preferentially in Tregs and, as a functional consequence, regulates Treg OXPHOS, which becomes more obvious in aged mice.

### Dj-1 regulates Treg maintenance via PDH

Having observed that DJ-1 regulates PDH activity via interacting with PDHB in nTregs and that DJ-1 depletion impairs nTreg proliferation and cellularity in aged mice, we further asked whether there is any causal link from the detected DJ-1-PDHB interaction to the observed Treg phenotypes. We therefore inhibited PDH activity to measure any potential effect on the proliferation and cell survival of WT nTregs sorted from young mice. Although we observed nTreg phenotypes in aged mice, we speculated the effects of PDH inhibition should already be visible in young mice as a specific PDH inhibitor should have a strong inhibition on PDH activity while DJ-1 only partially inhibits PDH activity. Compared with the vehicle treatment, the usage of a PDH inhibitor, CPI-613 (EC50 concentration around 200 uM in many tested tumor cell lines ^63^), essentially impaired proliferation, as assessed by a cell tracking dye (CellTracer Violet, CTV) and slightly compromised cell survival of nTregs following anti-CD3/-CD28/IL2 stimulation for 3 days (**Fig. 7a**). But the in vitro Treg lineage stability during the stimulation period of 3 days was not impaired by PDH inhibition (**Fig. 7a**). Since DJ-1 deficiency impaired SRC/maximum respiration in Tregs (**Fig. 6b**), we also inhibited PDH to evaluate whether SRC is affected in Tregs to further confirm whether DJ-1 regulates SRC via regulating PDH. As demonstrated, inhibiting PDH decreased SRC and maximum respiration in Tregs (**Fig. 6b**), indicating that DJ-1 regulates SRC via PDH.

**Figure 7.**
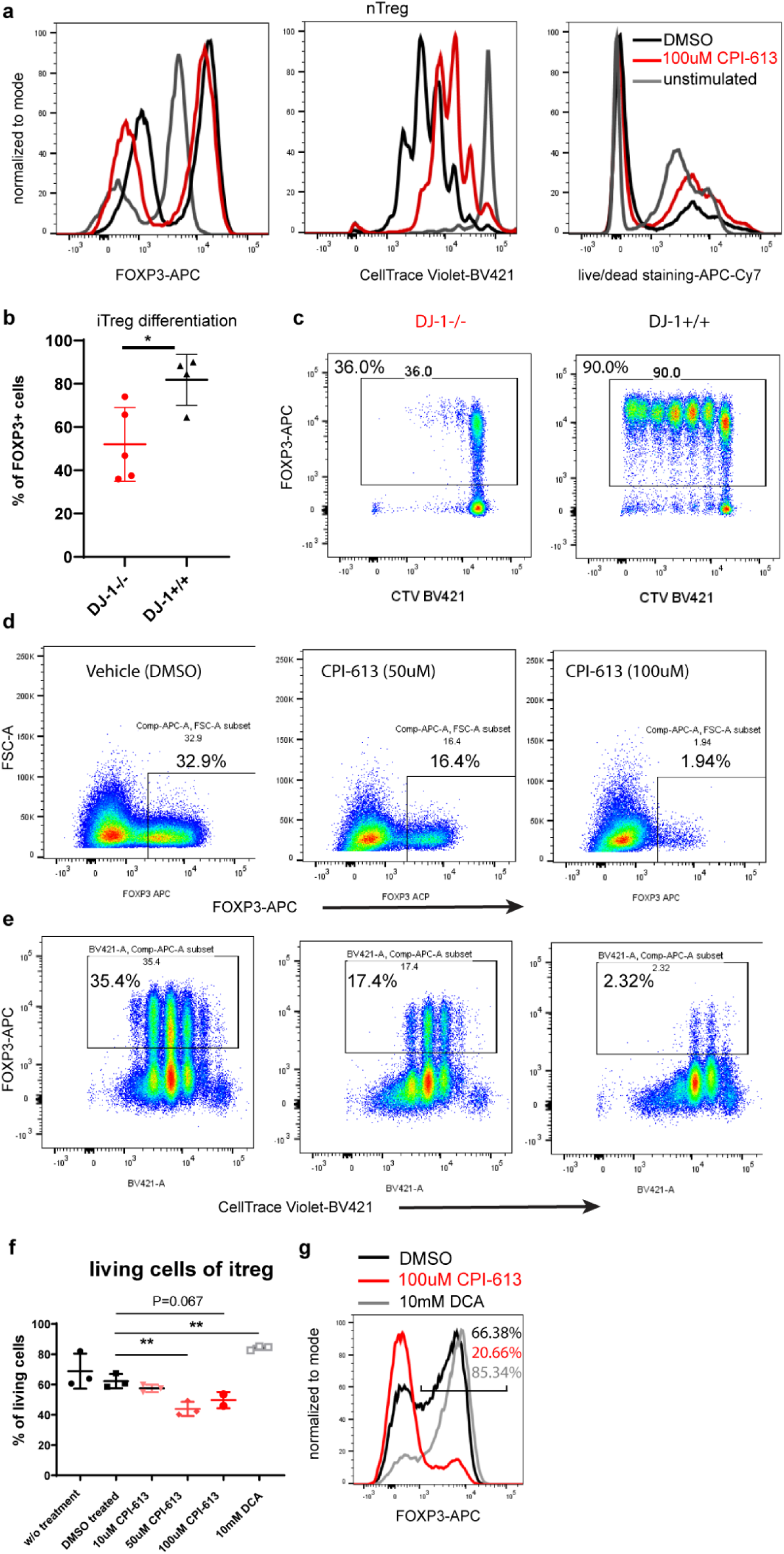
*Dj-1* depletion inhibits nTreg proliferation and iTreg differentiation via PDH. **a**, Representative histogram overlay of FOXP3 expression, cell proliferation (as labelled by celltrace violet [CTV]) or cell survival as measured by live/dead staining among CD4+ cells differentiated from Th0 cells under the iTreg differentiation condition stimulated for 3 days treated with PDH inhibitor (CPI-613) or control vehicle or without stimulation. **b**, Percentages of FOXP3 expression among DJ-1 KO or WT CD4+ cells differentiated from Th0 cells under the iTreg differentiation condition stimulated for 3 days. **c**, Representative flow-cytometry plots of FOXP3 and CTV staining among DJ-1 KO or WT CD4 cells differentiated from Th0 cells under the iTreg differentiation stimulated condition for 3 days. **d**, Representative flow-cytometry plots of FOXP3 expression among total WT CD4 T cells differentiated from Th0 cells treated with different doses of CPI-613 or control vehicle. **e**, Representative flow-cytoemtry plots of FOXP3 and CTV staining among total WT CD4 T cells differentiated from Th0 cells treated with different doses of CPI-613 or control vehicle. **f**, Percentage of living cells under the iTreg differentiation condition for 3 days with different doses of CPI-613 or DCA or control vehicle. **g**, Representative histogram overlay of FOXP3 expression differentiated from Th0 cells treated with CPI-613 or DCA or control vehicle. The percentages of FOXP3 expressing cells among gated living CD4 T cells were marked. Results represent four (**a, d, e**), three (**f, g**) and two (**b, c**) independent experiments. Data are mean± s.d. The P-values are determined by a two-tailed Student’s *t*-test. ns or unlabeled, not significant, *P<=0.05, **P<=0.01 and ***P<=0.001.

In line with the work done by others ^28^, *Dj-1* depletion also impaired differentiation and proliferation of iTregs (**Fig. 7b, c**). We therefore assessed whether these effects occur via PDH. Notably, inhibiting PDH activity in naïve T cells with CPI-613 significantly weakened the differentiation toward iTreg in a dose-dependent manner (**Fig. 7d**). Inhibiting PDH activity also impaired proliferation and cell survival although the administration dose of CPI-613 (ranging from 50 uM to 100 uM) was lower than the EC50 concentration (**Fig. 7e, f**). The effect on proliferation was already noticeable from the CPI-613 treatment with a concentration as low as 50 uM (**Fig. 7e**). A reduction in iTreg survival was also observed following *Dj-1* depletion ^28^. Since PDH activity is inhibited by PDH kinases, we assumed that inhibition of PDH kinases should generate opposite effects to PDH blockade. As speculated, treating naïve T cells with 10 mM of DCA (dichloroacetate, a PDH kinase inhibitor) relative to the control, we indeed observed an enhanced differentiation of iTregs (**Fig. 7g**), in line with the observations made elsewhere ^64^. In short, PDH inhibition phenocopied *Dj-1* depletion in nTregs and iTregs, indicating that DJ-1 maintains total Treg cellularity via regulating PDH activity in consideration of our observation that DJ-1 regulates PDH activity.

## Discussion

Here we uncovered a novel critical link between DJ-1 and PDHB that affects T cell immunity with a preferential impact on Tregs relative to the other CD4+ T cells (i.e., Tconv). We found that *Dj-1* depletion impaired Treg proliferation and the maintenance of Treg cellularity in older adult mice only. Another key observation of our study was that the apparent dysregulation in nTreg homeostasis of aged mice had a functional consequence by aggravating the symptom score during the remission phase of the induced autoimmunity disease model EAE. Pyruvate dehydrogenase (PDH) is a multiunit gatekeeper enzyme controlling the entry of glucose-derived pyruvate into the tricarboxylic acid cycle (TCA), a vital pathway providing cellular energy. Mitochondrial dysfunction in general ^2,65^, defects in PDH enzyme subunits ^66^, deficiency of critical cofactors of the PDH enzyme (e.g., thiamine ^67^) or deleterious mutations in pyruvate transporters ^68,69^ contribute not only to a variety of metabolic diseases ^5^, but also to several major neurological diseases, such as PD. At the same time, particular homozygous mutations in *DJ-1* cause 100% penetration of early-onset familial PD, which so far has been attributed mainly to the loss of anti-oxidative function in a cell-autonomous manner in neurons ^9-11^. Moreover, *Dj-1* has been shown to regulate mitochondrial respiratory functions and mitochondrial integrity ^12,13,70^. The effects of DJ-1 on mitochondrial functions might be attributable to cell type- and tissue microenvironment-specific regulations that eventually contribute to a complex disease phenotype. Although the previous functional and genetic studies described above have already suggested a functional convergence of DJ-1 and the PDH subunits leading to similarities in the phenotypic outcome, it remained elusive until now whether and how these two key cellular regulators interact. In line with the previously reported independent observations on DJ-1 and PDH functions already indicating convergent phenotypes, our unbiased proteomic data together with both, in vitro and in vivo analyses on the molecular and cellular level have revealed a physical and functional link between DJ-1 and PDH, which has a key impact on mitochondrial bioenergetic functions in nTregs, a crucial subset of CD4 T cells. Furthermore, we also showed that PDH inhibition phenocopied the effects of DJ-1 deletion in Tregs. Thus, our findings add several new aspects to the understanding of the DJ-1/PARK7-based monogenic defect, which leads to early onset PD with full penetrance: i) the finding of significant immune dysregulation affecting T cell immunity beyond the cell-autonomous dysregulation already described in neurons; ii) the obvious age-dependency of the observed immune phenotypes; and iii) the layer of critical interaction of major intracellular metabolic pathways at the crossroads of mitochondrial functions and the TCA cycle.

While loss-of-function defects or inhibition of DJ-1 and PDH lead to partially overlapping and convergent cellular and disease phenotypes, no direct interaction of these two key cellular regulators has been described so far. Our results indicated that DJ-1 regulates PDH activity and OXPHOS in Tregs in a novel and unexpected way. Through mass spectrometry analysis followed by anti-DJ-1 co-immunoprecipitation in unstimulated and stimulated human Tregs and Tconv, we detected that PDHB was the most prominent protein with binding preference for DJ-1. Furthermore, we found a much higher abundance of PDHB protein bound to DJ-1 in unstimulated cells than stimulated ones. The binding between DJ-1 and PDHB indeed regulated the enzymatic activity of PDH via modulating phosphorylation levels of PDHA mainly in Tregs and in resting Tconv. Due to the preferential interaction between DJ-1 and PDHB in Tregs, the loss of DJ-1 reduced OXPHOS capacity in Tregs as a functional consequence, especially in aged mice. These findings are in line with previously reported data where it has been shown that Tregs rely more on OXPHOS than Tconv ^62,71^. Furthermore, the biased effect on PDH activity in unstimulated T cells can further be explained by the fact that T cells switch their metabolic pathway mode from OXPHOS to glycolysis upon activation ^57^. Thus, our study has identified a novel and hitherto unknown protein-protein interaction between DJ-1 and PDH that connects two major cellular pathways, mitochondrial functions with DJ-1 as a crucial player and the TCA cycle with PDH as a key enzyme.

An appropriate homeostatic regulation of Treg activity and function is crucial for a balanced immune response during the physiological aging process^30^. Our study demonstrated that DJ-1 depletion impairs nTreg proliferation and cellularity in older adult mice only. Since OXPHOS is critical for the development and long-term survival of Tregs ^71^, it is plausible that the compromised spared respiratory capability, which we have demonstrated in aged mice devoid of *Dj-1*, contributed to the impaired Treg accumulation. The spared or maximum respiration is often not necessary under homeostatic and disease-free conditions, but is required under cellular stress or activation conditions such as in low-grade inflammation in the elderly, a state that is also known as ‘inflammaging’ ^72^. In line with this notion, age-dependent mitochondrial dysfunction has been observed in skeletal muscle of *Dj-1*-depleted mice ^13^. We further elucidated the underlying mechanisms and found that the impairment of nTreg proliferation and cellularity occurred via PDHB, as blockade of the PDH enzymatic activity phenocopied the effects of *Dj-1* deletion on nTreg cellularity and OXPHOS capacity. The functional consequences of PDH inhibition through *Dj-1* deletion on Treg cellularity appear to be compensated by other pathways during young age ^73^. However, these effects on Treg proliferation and cellularity cannot be balanced anymore in old *Dj-1*-deficient mice beyond a critical age and tipping-point ^74^. In our study we have applied an inhibitor of PDH that phenocopied the effects of *Dj-1* depletion in purified nTregs. In line with our findings, the administration of the same PDH inhibitor in preclinical trials induced systemic inflammatory responses as the primary and only side effect, which is indicative of systemic immune dysregulation ^75^. Thus, our data provide strong evidence that *Dj-1* deletion compromises nTreg cellularity and accumulation in aged mice only, via a previously unrecognized interaction of DJ-1 with PDHB.

The depletion of *Dj-1* has previously already been linked to an impaired differentiation of iTregs from naïve CD4 T cells in young adult mice ^28^. Our results confirm that *Dj-1* depletion impairs iTreg differentiation. However, the study by Singh and coworkers ^28^ provided no clear-cut underlying molecular mechanism of how iTreg differentiation is impaired in *Dj-1 KO* mice. Our results showed that inhibiting PDH also impaired differentiation, proliferation and survival of iTregs in an identical manner as it was observed with *Dj-1* deletion. Since OXPHOS is required for activation of naïve T cells ^76^, it is not surprising that PDH blockade impaired the downstream TCA-driven OXPHOS and consequently compromised differentiation of naïve T cells towards iTregs. Interestingly, depletion of *Dj-1* did not impede Treg suppressor function, which is indicative that OXPHOS might be more vital to the long-term survival rather than the suppressive function of Tregs. Together, these results demonstrate that DJ-1 regulates overall Treg cellularity via modulating PDH enzyme activity.

Since Tregs are critical during the stable and remission phases of various autoimmune diseases ^48^, we hypothesized that reduced Treg cellularity, as observed in our study after *Dj-1* depletion, should worsen disease symptoms. As a critical role of Tregs has especially been demonstrated in EAE ^77^, a murine model for the human autoimmune disease multiple sclerosis, we decided to analyze the role of DJ-1 further in this particular autoimmune disease model. Indeed, our data showed that depletion of *Dj-1* deteriorated symptoms during the remission-phase of EAE in old mice as compared to age- and gender-matched WT mice. As no difference in Treg frequency was observed between young *Dj-1 KO* and WT mice, we also did not observe a difference in the EAE scores between the corresponding young adult groups. Therefore, *Dj-1* depletion impaired Treg cellularity and consequently Treg pathophysiological functions in aged mice. We thus concluded that *Dj-1* depletion enhances autoimmune responses in a well-characterized disease model in an age-dependent way.

Although PDH enzyme activity is tightly regulated by several PDH kinases and phosphatases, our findings on DJ-1 promoting PDH activity add a new dimension to the regulation of the PDH complex, thus enabling a better understanding of the highly organized PDH supercomplex system ^78^. Although we cannot exclude potential effects of other cell types on the degree of accumulation of total Tregs in mice with a constitutional deletion of *Dj-1* used in this study, our work underscores that the DJ-1-PDHB interaction could be used as a potential pharmaceutical target to interfere with various complex diseases involving dysregulation of Treg frequency in the aged population. Uncovering this unexpected critical physical and functional interaction of the familial PD gene DJ-1 on PDH activity in Tregs suggests that other genes involved in neurodegenerative diseases and regulating mitochondrial respiratory functions might be among potential candidates that steer the activity of PDH or other TCA enzymes. In line with our discoveries, another PD familial gene, PARK2/PARKIN, regulates the important glycolysis rate-limiting enzyme pyruvate kinase M2 in tumor cell lines ^79^. However, pyruvate kinase M2 is not expressed in human CD4 T cells. Our data demonstrate that DJ-1 counteracts the role of the PDH kinases, e.g., PDK1 on PDH ^64^, mainly in the specific CD4 T-cell subset of Tregs. Future experiments are needed to determine whether the observed DJ-1-PDHB interaction also holds true in dopaminergic neurons and accordingly contributes to the death of such neurons, which essentially causes the motor dysfunction of PD ^16^.

## Materials and Methods

### Human T cell-related experiments

#### Human T cell isolation

The buffy coats or leukopaks from healthy donors used for primary T cell culture and analysis were provided by the Luxembourg Red Cross. Human primary natural Tregs and Teffs (CD4 effector T cells) were derived respectively from sorted CD4^+^CD25^high^CD127^low^ and CD4^+^CD25^-^ T cells of peripheral blood mononuclear cells (PBMC) on BD FACSAria™ III cell sorter. The PBMCs were isolated by using Ficol-Paque plus (17-1440-03, GE Healthcare) or Lympoprep (07801, StemCell) followed by magnetic separation with anti-human CD4 microbeads (130-045-101, Miltenyi Biotec), as described by the manufacturers, before being stained with mouse monoclonal [RPA-T4] anti-human CD4 FITC (555346, BD Biosciences) (dilution 1:20), mouse monoclonal [M-A251] anti-human CD25 APC (555434, BD Biosciences) (dilution 1:20) and mouse monoclonal [HIL-7R-M21] anti-human CD127 V450 (560823, BD Biosciences) (dilution 1:20) for sorting with BD FACSAria™ III.

#### Human T cell culturing

We followed the same protocol as descried in our previous work ^41^. Sorted T cells were cultured in IMDM (21980-032, Thermo Fisher Scientific) complete medium supplemented with 10% heat-inactivated (56°C, 45 min) Gibco® fetal bovine serum (FBS) (10500-064, Thermo Fisher Scientific), 1x Penicillin+Streptomycin (15070-063, Thermo Fisher Scientific), 1x MEM non-essential amino acids (M7145, Sigma-Aldrich) and 50 μM beta-mercaptoethanol (M7522, Sigma-Aldrich). All the human T cells were cultured in 37°C 7.5% CO2 incubators, unless specified. 100 U/ml recombinant human IL2 (known as Proleukin® in medication) (PZN 2238131, Novartis) was added daily to Treg (but not Teff) cell culture medium and the same amount of IL2 was added to Tregs unless otherwise stated. Every seven days, all T-cells were restimulated with irradiated Epstein–Barr virus (EBV preparation from VR-1492, ATCC)-transformed B-cells ^80^ (EBV-B cells), at a 1:1 ratio of T and EBV-B cells, to expand and maintain the T-cells. The EBV-B cells were irradiated in an RS2000 X-Ray Biological Irradiator (Rad Source Technologies) for 30 min at a rate of 2.80 Gy/min. The T-cells were regularly characterized by co-staining CD4, CD25, FOXP3 and Helios protein levels by flow cytometry (**Fig. EV1a, b**). When the primary Tregs were older than 6 weeks or the expression level of FOXP3 or Helios or the cellular viability was apparently decreased, the cells were discarded and new T cells were isolated from different healthy donors. The antibodies used for cell quality assessment were: mouse monoclonal [RPA-T4] anti-human CD4 BUV395 (564724, BD Biosciences) (dilution 1:100), mouse monoclonal [M-A251] anti-human CD25 FITC (555431, BD Biosciences) (dilution 1:100), mouse monoclonal [22F6] anti-human Helios Pacific blue (13721, BioLegend) (dilution 1:100), mouse monoclonal [206D] anti-human FOXP3 Alexa Fluor 647 (320119, BioLegend) (dilution 1:10). The LIVE/DEAD® Fixable Near-IR Dead Cell Stain (L10119, Thermo Fisher Scientific)(dilution 1:500) was used to distinguish living cells from dead cells. In certain cases, we directly compared the markers of our isolated human Tregs with TregThu, a golden standard isolated from our previous work ^80^. The procedure for the staining of extracellular and intracellular markers is described below.

#### Gene knockdown in human T cells

Targeted gene expression was knocked-down by using P3 Primary Cell 4D-Nucleofector X Kit L (V4XP-3024, Lonza) with 100 µl P3 primary cell solution and 100 pmol of the corresponding si_RNAs (resuspended in 10 μl RNAse-free H2O): si_Non-Specific (si_NS or si_CTL) (sc-37007, Santa Cruz) and si_PARK7/_DJ-1 (SI00301091, *AATG GAGG TCAT TACA CCTA C*, Qiagen). Amaxa 4D-Nucleofector™ X System (Lonza) was used for the experiments with the manufacturer’s recommended program for stimulated human primary T cells. Following the siRNA transfection procedure, T cells were transferred into 12-well plates with pre-warmed complete medium supplemented with 100 U/ml IL-2 for Tregs and kept at 37°C for 1 day. They were then stimulated with Dynabeads Human T-Activator CD3/CD28 (11131D, Thermo Fisher Scientific) (ratio of cells and beads: 1:1) or with ImmunoCult Human T Cell Activator (10991, STEMCELL Technologies) (25 μl/ml) in 24-well plates for 1 or 2 days depending on the corresponding experiments, with or without additional recombinant IL-2 for Tregs and Teffs, respectively.

#### RNA extraction and cDNA synthesis

RNA was extracted as previously described ^41^. RNA samples for standard and quantitative PCR were prepared by using the RNeasy Mini Kit (74106, Qiagen) starting with lysing the cells with RLT buffer supplemented with 1% beta-mercaptoethanol (63689, Sigma-Aldrich), following the manufacturer’s instructions and including the digestion of genomic DNA with DNAse I (79254, Qiagen). The RNA concentration was measured with a NanoDrop 2000c Spectrophotometer (Thermo Fisher Scientific) followed by a quality check of the RNA integrity number (RIN). For RIN assessment, the Agilent RNA 6000 Nano kit (5067-1511, Agilent) was used together with the Agilent 2100 Bioanalyzer Automated Analysis System (Agilent) according to the manufacturer’s protocol.

A maximum of 500 ng RNA was used for cDNA synthesis. A master mix for the first step was prepared with 0.5 μl of 50 μM Oligo(dT)_20_ primers (18418020, Thermo Fisher Scientific), 0.5 μl of 0.09 units/μl Random Primers (48190-011, Thermo Fisher Scientific), 1 μl of 10 mM dNTP mix (R0192, Thermo Fisher Scientific) and RNAse-free water made up to a final volume of 13 μl in 0.2 ml PCR Tube Strips (732-0098, Eppendorf). The tubes were transferred into a Professional Standard Gradient 96 Thermocycler (Biometra) for 5 min at 65°C and 2 min at 4°C. After the first step, the reaction was supplemented with 40 units RNaseOUT™ Recombinant Ribonuclease Inhibitor (10777019, Thermo Fisher Scientific), 200 units SuperScript™ III Reverse Transcriptase (18080-044, Thermo Fisher Scientific) and dithiothreitol (DTT) (70726, Thermo Fisher Scientific) to give a final concentration of 5 mM in a total reaction volume of 20 μl. The PCR tubes were returned to the thermocycler at 50°C for 60 min, 70°C for 15 min and 4°C until further usage.

#### Standard polymerase chain reaction (ST-PCR) and realtime quantitative PCR (qPCR)

The DreamTaq Green PCR Master Mix (K1081, Thermo Fisher Scientific) was used as a base for the ST-PCR. Forward and reverse primers together with cDNA and RNAse-free water were added. PCR was performed in a Professional Standard Gradient 96 Thermocycler (Biometra). Following the amplification, the samples and MassRuler DNA Ladder Mix (SM0403, Thermo Fisher Scientific) were loaded onto 2% agarose (A9539, Sigma-Aldrich) gel with ethidium bromide (E1510, Sigma-Aldrich) or SYBR Safe (S33102, Thermo Fisher Scientific, dilution 1:10 000) and the gel was run for 1 hr at 120 V in TAE buffer. Images of the bands were taken with the G:Box gel doc system (Syngene).

Real-time quantitative PCR (RT-PCR/qPCR) was performed by using LightCycler 480 SYBR Green I Master Mix (04707516001, Roche), supplemented with cDNA and primers in a reaction volume of 10 μl. LightCycler 480 Multi-well White Plates (04729749 001, Roche) with 384 wells and LC 480 Sealing Foil (04729757001, Roche) were used in these experiments. The reaction was undertaken on a LightCycler 480 (384) platform (LightCycler 480 (384), Roche). The results were analysed with LightCycler 480 SW 1.5 software. The annealing temperature of 55°C was set for the various genes unless stated. The primers used for human qPCR were: PARK7 (QT00055811, Qiagen), RPS9 (QT00233989, Qiagen) or GAPDH (QT00079247, Qiagen) as reference genes.

#### Human suppression assay

Human Teff cells were stained with 1 μM CFSE (carboxyfluorescein succinimidyl ester, C34554, Thermo Fisher Scientific) dye in PBS for 2min 45sec in the dark at room temperature. The reaction was stopped with cold FBS (2min, dark, 4°C) before the cells were washed and recultured in complete IMDM medium. Tregs were transfected with scrambled control or gene-specific siRNA for 1 day before being cocultured with Teffs and EBV-B cells. Serial dilution of Tregs was performed in a 96-well plate with a starting cell number of 6E4/well. 6E4 CFSE-stained Teff cells were distributed in each well with various amounts of Treg cells and stimulated with 6E4 irradiated EBV B-cells (irradiation conditions were the same as for the cell culture). As positive or negative proliferation controls, the same number of CSFE-stained Teff cells were cultured without Tregs, with and without irradiated EBV B-cells. The cell mixtures were cultured at 37°C for 5 days and FACS analysis was carried out to determine the proliferation capacity of Teff cells by additionally staining with CD4 APC ab and the LIVE/DEAD® Fixable Near-IR Dead Cell Stain reagent.

#### Co-immunoprecipitation (Co-IP)

The same number (around 1E7 T cells) of Tregs and Teffs were either left unstimulated or were stimulated for 5 hrs with Dynabeads Human T-Activator CD3/CD28 for cell expansion and activation (11131D, Thermo Fisher Scientific) (ratio of 1:1). After 5 hrs, cell pellets were collected, lysed in homogenization buffer with 0.2% Triton X100, 100 mM NaCl, 2 mM EDTA and 50 mM Tris (pH=8.0). Lysates were pre-cleared with 10 ul washed Dynabeads Protein G (10003D, Thermo Fisher Scientific) in 10 ul lysis buffer for 1 hr at room temperature and the beads were then removed. The total proteins in each sample were quantified by standard Bradford assay (5000006, Bio-Rad) in a microplate Cytation 5 cell imaging multi-mode reader (1509096, BioTek Instruments GmbH). The lysates were diluted to a 3 µg/µl protein concentration with protease inhibitor cocktail (PIC) at a dilution of 1:100 to give the final volume. The samples were then evenly distributed into two parts. One part of the sample was immune-precipitated with rabbit monoclonal [EP2815Y] anti-human PARK7/DJ-1 antibody (ab76008, Abcam) and another part was immune-precipitated with normal rabbit IgG (sc-2027, Santa Cruz) as a negative control, for 1 hr at room temperature (for both target and control antibodies, the dilution was 1:10) before the addition of Dynabeads (20 µl beads for 7 µl of antibody) overnight at 4°C with gentle rocking. The beads were washed 3x in lysis buffer and sequentially 2x in lysis buffer without detergent. The DJ-1 ab pull-out complexes were then eluted from the beads by using 0.1% RapiGest (186001861, Waters) in 50 mM ammonium bicarbonate (A6141, Sigma-Aldrich) solution with 5 mM DTT (43819, Sigma-Aldrich) at 60°C for 1 hr, dried in a SpeedVac Concentrator at 4°C (Thermo Fisher Scientific) and resuspended in a small volume of lysis buffer (for Western blot) or in ultra-pure H2O with 1% acetonitrile (AE70.2, Carl-Roth) (for mass spectrometry analysis).

#### Western Blotting (WB)

Protein samples were mixed 1:3 with 4x Laemmli sample buffer (NP0007, Life Technologies) containing 50 mM dithiothreitol (DTT), heated for 5min at 95°C, cooled on ice and loaded on gradient PAGEr™ Gold Precast Gels (58505, Lonza). ProSieve™ EX Running (200307, Westburg) and ProSieve™ EX Transfer (200309, Westburg) buffers were used in the assay according to manufacturer’s recommendations and both PageRuler™ Prestained Protein Ladder (26616, Thermo Fisher Scientific) and Precision Plus Protein™ All Blue Prestained Protein Standards (Bio-rad, 1610373) were used to determine the size of the protein bands. Proteins were transferred to a methanol (34860, Sigma-Aldrich)-activated (5min, room temperature) Immun-Blot PVDF membrane (162-0177, Bio-Rad) and then blocked with 5% milk (70166, Sigma-Aldrich) in PBS (14190169, Thermo Fisher Scientific) with 0.2% Tween-20 (P7949, Sigma-Aldrich) for 1 hr at room temperature with gentle shaking. All primary antibodies were diluted in the blocking solution and incubated overnight at 4°C. The membrane was washed (3×10min) with PBS-Tween-20 (0.2%) before and after incubation with secondary antibodies. Chemiluminescence-based Amersham ECL Prime Western Blotting Detection Reagent (RPN2232, VWR/GE-Healthcare) was employed to detect the bands, which were visualized in the Westburg Odyssey FC Imaging System (Li-COR). If needed, the membrane was stripped with Restore Western Blot Stripping Buffer (21059, Thermo Fisher Scientific) for 30min at room temperature, washed, blocked and blotted again. The main antibodies used for Western blotting were: rabbit monoclonal [EP2815Y] anti-human PARK7/DJ-1 antibody (ab76008, Abcam) (dilution 1:2000), rabbit Anti-Pyruvate Dehydrogenase E1-alpha subunit [EPR11098] (ab168379, Abcam) (dilution 1:3000), rabbit anti-human Pyruvate Dehydrogenase E1 beta subunit (PA5-31519, Thermo Fisher Scientific) (dilution 1:1000), rabbit anti-human Pyruvate Dehydrogenase E1-alpha subunit (pSer293) (ab92696, Abcam)(dilution 1:300), rabbit anti-human phospho PHE1-A type I (pSer300) (ABS194, Merck Millipore)(dilution 1:1000), rabbit monoclonal [6C5] anti-human GAPDH (sc-32233, Santa Cruz) (dilution 1:1000). A goat anti-rabbit IgG (H+L)-HRP conjugated antibody (172-1019, Bio-Rad) (dilution 1:5000) was used as the secondary antibody.

For the assessment of the phosphorylation of PDHE1A, DJ-1/PARK7 were knocked-down in 5E6 Treg cells or in 5E6 Teff cells, according to the manufacturer’s protocol for 24 hrs (P3 Primary Cell 4D-Nucleofector® X Kit, Lonza); see the section ‘Gene Knockdown’). The siRNA transfected cells were then stimulated with ImmunoCult Human CD3/CD28 T Cell Activator (25 µl/ml) (Stemcell Technologies, France) for 5, 24 or 48 hrs. Tregs were supplemented with IL-2 (100 U/ml) daily for the duration of the experiment. Proteins were quantified, separated, transferred, blocked and stained by means of the same approaches as those described above. Detection was carried out on an ECL Chemocam Imager (INTAS). For antibody information, please refer the paragraph above. If needed, the contrast and brightness of the whole image was adjusted by using the *ImageJ* (https://imagej.nih.gov/ij/index.html) free software.

#### Mass spectometry (MS)

The quality of the samples prepared for MS analysis was checked by Western Blot by loading a small amount of the re-suspended IP elution products onto acrylamide gel. Only if the bait protein (PARK7/DJ-1) was detected on the blotted membrane, was the IP considered successful and only these samples were analysed by mass spectrometry in the proteomics facility (Institute for Systems Biology, Seattle). Eluted proteins were reduced with 5 mM dithiothreitol (DTT), alkylated with 25 mM iodoacetamide and digested with Lys-C for 3 hrs at 37°C (11420429001, Promega) (dilution 1:200) followed by an overnight digestion with Trypsin (V5113, Promega) (dilution 1:25) at 37°C. Peptides were acidified with formic acid (33015, Sigma-Aldrich) to stop digestion and remove RapiGest. Peptides were purified by using C18 reversed-phase chromatography followed by hydrophilic interaction chromatography (HILIC, Nest Group). We separated purified peptides by online nanoscale HPLC (Agilent) with a C18 reversed-phase column packed 14 cm (ReproSil-Pur C18-AQ 3 µm; Dr. Maisch Products) over an increasing gradient of 3-35% Buffer B (100% acetonitrile, 0.1% formic acid) for 90 min at a flow rate of 300 nl/min. Eluted peptides were analyzed with an Orbitrap Elite mass spectrometer (Thermo Fisher Scientific) operated in a data-dependent mode. The top 15 most intense peptides per MS1 survey scan were selected for further MS2 fragmentation by collision-induced dissociation (CID). We performed MS1 survey scans in the Orbitrap at a resolution of 240,000 at m/z 400 with charge state rejection enabled, whereas CID MS2 was done in the dual linear ion trap with a minimum signal of 1000. We set dynamic exclusion to 30 sec. We employed Maxquant (v1.5.2.8) to analyze raw output data files ^81^. Peptides were compared with the UniProt human database (02-2015 release). We imposed a strict 1% FDR cut-off to a reverse sequence database. The MaxLFQ algorithm was used for label-free quantification ^81^. Microsoft Excel and Perseus (v1.5.0.15) ^82^ were utilized for data processing. We removed contaminants, decoys and single peptide identifications for further analysis. Zero values in the remaining data were substituted by imputation. The results were pre-processed and certain criteria were used to validate previously known PARK7/DJ-1 binding partners and to predict and validate the as yet unknown ones. We selected potential novel binding partners by using a combination of the following criteria: (1) the average mRNA expression levels of the corresponding transcripts in Tregs over the first 6 hrs following TCR stimulation should be higher than 250 (the average expression level of FOXP3 in human natural Tregs from the time-series transcriptomic datasets is around 250); (2) the LFQ value of MS-identified proteins bound to DJ-1 antibody should be at least 2.5-fold higher than those bound to the IgG control in human primary T cells; (3) the fold changes in the LFQ value of the bound proteins between stimulated and unstimulated T cells for all the three analysed conditions (Teffs, Tregs with or without IL-2 addition) should be equal to or higher than 2; (4) a similar change trend (increase or decrease) in binding affinity between stimulated and unstimulated samples should be consistent for all the three analysed conditions; (5) the LFQ value of the bound proteins in the eluates of Tregs, with or without IL2 addition, should be higher than the 25^th^ percentile of all the detected binding proteins in the given unstimulated or stimulated samples.

The known DJ-1 binding partners in human cells were downloaded from the NCBI database. Independent of the mRNA expression levels, the binding partners were considered confirmed when the following two criteria were met: (1) the eluted protein LFQ value was at least 2-fold higher than that in the corresponding IgG controls of either human T cells or the Jurkat cell line (ACC282, DSMZ GmbH); (2) the fold enrichment (stimulated vs. unstimulated) from at least 2 out of the 4 measured conditions (Teffs, Jurkat cells, Tregs with or without IL2 addition) was similar.

#### Flow cytometry (FACS) and ImageStream for human cells

Once the cells had been collected and the medium removed, the extracellular proteins of the cells were stained in FACS buffer (PBS (14190169, Thermo Fisher Scientific) containing 2% heat-inactivated FBS (10500-064, Thermo Fisher Scientific) with the cell surface antibodies, the concentration of which was first optimized, for 30 min at 4°C in the dark. Intracellular staining was performed by using the FOXP3 staining kit (421403, BioLegend) following the manufacturer’s instructions. Stained cells were analysed with BD LSRFortessa™. We excluded dead cells from the further analysis by using LIVE/DEAD® Fixable Near-IR Dead Cell Stain (L10119, Thermo Fisher Scientific) (dilution 1:500). Various combinations of the following antibodies were used for FACS analysis: mouse monoclonal [RPA-T4] anti-human CD4 APC (561840, BD Biosciences) (dilution 1:200), mouse monoclonal [RPA-T4] anti-human CD4 FITC (555346, BD Biosciences) (dilution 1:200), mouse monoclonal [G44-26] anti-human CD4 V450 (561292, BD Biosciences) (dilution 1:200), mouse monoclonal [RPA-T4] anti-human CD4 BUV395 (564724, BD Biosciences) (dilution 1:200), mouse monoclonal [L200] anti-human CD4 PE-Cy7 (560644, BD Biosciences) (dilution 1:200), mouse monoclonal [M-A251] anti-human CD25 APC (555434, BD Biosciences) (dilution 1:200), mouse monoclonal [M-A251] anti-human CD25 FITC (555431, BD Biosciences) (dilution 1:200), mouse monoclonal [M-A251] anti-human CD25 V450 (560356, BD Biosciences) (dilution 1:200), mouse monoclonal [HIL-7R-M21] anti-human CD127 V450 (560823, BD Biosciences) (dilution 1:200), mouse monoclonal [50G10] anti-human GARP (221 011, Synaptic Systems GmbH) (dilution 1:100), mouse monoclonal [B56] anti-human Ki-67 V450 (561281, BD Biosciences) (dilution 1:100), mouse monoclonal [206D] anti-human FOXP3 Alexa Fluor 647 (320119, BioLegend) (dilution 1:10), mouse monoclonal IgG1 (50-167-013, BioLegend) (dilution 1:10), mouse monoclonal [22F6] anti-human Helios Pacific blue (137210, BioLegend) (dilution 1:100), CellTrace CFSE Cell Proliferation Kit (C34554, Thermo Fisher Scientific) (dilution 1:5000). For staining GARP expression, anti-mouse IgG3 (biotin) (553401, BD) and Streptavidin-Phycoerythrin (PE, 349023, BD Biosciences) were used as secondary antibodies. The FCS files from human-related experiments were analysed by FlowJo 7.6.5 or FlowJo v10 (Tree Star).

For ImageStream experiments, Tregs and Teffs were stimulated and stained as described above, but with staining panels other than those for the FACS analysis. We used the following staining panels: LIVE/DEAD® Fixable Near-IR Dead Cell Stain (L10119, Thermo fisher Scientific) (dilution 1:500), mouse monoclonal [RPA-T4] anti-human CD4 V450 (560346, BD Biosciences) (dilution 1:100), rabbit monoclonal [EP2815Y] anti-human PARK7/DJ-1 Alexa Fluor 488 (ab203989, Abcam) (dilution 1:50), rabbit monoclonal [EPR11097(B)] anti-human PDHB Alexa Fluor 594 (ab211838, Abcam) (dilution 1:100), including strict negative controls (rabbit monoclonal [EPR25A] IgG Alexa Fluor 488 (ab199091, Abcam) (dilution 1:50) and rabbit monoclonal [EPR25A] IgG Alexa Fluor 594 (ab208568, Abcam) (dilution 1:100). The samples were acquired on an ImageStream®^X^ Mark II Imaging Flow Cytometer (Amnis, EMD Millipore) at 60x magnification. The results were analysed by using IDEAS 6.2 software (Amnis) and the Similarity score between PDHB and DJ-1 was determined by means of a similarity dilate algorithm often used for the nuclear translocation quantification of a transcription factor. We utilized the PDHB-positive area as the targeted location and then calculated to what degree DJ-1 was translocated to the PDHB-positive areas. Only cells gated as living focused singlets were included in the analysis.

#### PDH enzymatic assay

Various genes were knocked-down for the enzymatic assay, as described in the previous sections. The same number (1E6) of Treg and/or Teff cells were either left unstimulated or stimulated for 5 hrs with Dynabeads Human T-Activator anti-CD3/-CD28 (11131D, Thermo Fisher Scientific; the ratio between beads and T cells is 1:1) with (Tregs) or without IL-2 (Teffs). Cell pellets were then collected and either lysed in lysis buffer or stored at -80°C until further analysis. Protein concentrations were determined by using the Bradford assay and the same amount of protein was used for each sample within one experiment. PDH activity was determined by means of the colorimetric PDH Activity Assay Kit (MAK183, Sigma-Aldrich) following the manufacturer’s recommendations in Nunc MicroWell 96-well microplates (167008, Thermo Fisher Scientific) in an Infinite 200 Pro plate reader (Tecan) measuring the absorbance at 450 nm every 10-30 seconds over a time course of 1-5 hrs. Various amounts of recombinant DJ-1 protein (ab51198, Abcam) were used (0-20 µg) to test the effect of recombinant human DJ-1 on PDH activity.

#### Nucleotide analysis

The gene knockdown protocol described above was used to prepare knockdown conditions in Treg cells, namely control si_ NS (sc-37007, Santa Cruz) and si_PARK7/DJ-1 (SI00301091, Qiagen) and two knockdown conditions in Teff cells, namely si_NS (sc-37007, Santa Cruz) and si_DJ-1 (SI00301091, Qiagen). On the following day, half of the samples were stimulated with Dynabeads Human T-Activator CD3/CD28 for cell activation (11131D, Thermo Fisher Scientific) for 1 day with IL2 (Tregs) or without IL-2 (Teffs) in 48-well plates in a total volume of 1ml complete IMDM medium+10% FBS. An aliquot of 0.5 ml cell suspension was transferred to a 2 ml Eppendorf tube containing 1.5 ml of 60% methanol, 0.85 % AMBIC, pH=7.4 at -60°C. The tubes were gently shaken before the cells were separated by centrifugation (3 min, 500 G, -10°C). The supernatant was discarded and the cell pellet was stored at -80°C until intracellular metabolite extraction. Intracellular metabolites were extracted by using 50% Methanol TE buffer, pH=7.0 at -20°C. Chloroform at - 20°C was used to improve the cell lysis. Cell lysates were taken from the freezer, resuspended in extraction fluid and chloroform and incubated on a shaking device at -20°C for 2 hrs. Next, cell debris were separated by centrifugation (10 min, 10 000 rpm, -10°C). The extract was filtered (0.22 µm CA) and stored at -80°C until analysis. The volume of extraction fluid and chloroform was adjusted to the cell number in the pellet (0.5 ml per 3E6 cells). Extracted nucleotides were separated on a SeQuant® ZIC®-HILIC 3.5µm, 100 Å 150 × 2.1 mm column (Merck Millipore) at 25 °C and 0.400 mL/min and analyzed on a Dionex UltiMate 3000 UHPLC Systems coupled to a Q Exactive™ Hybrid Quadrupole-Orbitrap™ Mass Spectrometer (Thermo Fisher Scientific). The separation of the nucleotides was carried out using a binary gradient with buffer A: water and B: 90 % acetonitrile, both containing 20 mM ammonium acetate with pH 7.5. The liquid chromatography started with a 10 min liner gradient of 90 – 50 % of B, followed by a second liner gradient of 50 – 20 % B for 1 min. The washing step was completed within 1 min by increasing buffer B to 90 %, followed by 8 min of column equilibration with initial condition. Full MS scans were acquired at a target value of 3e6 ions with resolution R = 70,000 over 3 mass ranges of 345.5 – 346.5; 425.5 – 426.5; 505.5 – 506.5 m/z.

### Mouse-related experiments

B6.129P2-Park7^Gt (XE726) Byg^/Mmucd mice were developed and characterized as described elsewhere ^46^. The *Dj-1*^*-/-*^ *(KO), Dj-1*^*+/-*^, *Dj-1*^*+/+*^ (WT) mice used in our experiments were gender- and age-matched siblings generated from heterozygous *Dj-1*^*+/-*^ breeding pairs. All mice were maintained in our SPF animal facility and all animal experimental procedures were performed following the approval of the Animal Welfare Society (AWS) of University of Luxembourg and Luxembourg Institute of Health.

#### FACS for murine T cells

The same number of cells from spleen or peripheral lymph nodes were first incubated with anti-mouse CD16/CD32 Fc blocker (553141, BD Biosciences) and then stained with various surface and intracellular antibodies. Cell numbers were determined by CASY (Innovatis AG). Cell surface markers were stained by rat anti-mouse CD4-BUV496 (564667, BD Biosciences) (dilution 1:200), rat anti-mouse CD25-PE-Cy7 (25-0251-82, eBioscience) (dilution 1:200), rat anti-mouse CD8-BUV805 (564920, BD Biosciences) (dilution 1:200), hamster Anti-Mouse PD-1-BV711 (744547, BD Biosciences) (dilution 1:100), rat anti-mouse GITR-BUV395 (740322, BD Biosciences) (dilution 1:100) and rat anti-mouse ICOS-FITC (11-9942-82, eBioscience) (dilution 1:100). Of note, all these antibodies might not be necessarily used in one staining panel due to the spectrum compatibility. For cytokine staining, cells were fixed and permeabilized with Cytofix/Cytoperm buffer (554714, BD Biosciences). Anti-IL-10 (505028, BioLegend) antibody was diluted in Perm/Wash buffer (554714, BD Biosciences). Intracellular staining for Foxp3 (17-5773-82, eBioscience) (dilution 1:100), CTLA4 (12-1522-83, eBioscience) (dilution 1:100), Helios (137220, BioLegend) (dilution 1:100) and Ki67 (48-5698-82, eBioscience) (dilution 1:100) was performed by using the Foxp3 staining kit (00-5523-00, eBioscience). Rat anti-mouse Ki67-BV605 (652413, BioLegend) (dilution 1:100) was used in the thymic analysis. Samples were measured on a BD LSRFortessa™ and data were analysed with FlowJo (v10, Tree Star).

#### Murine Treg suppression assay

Spleens and lymph nodes were minced through a 70 µM-pore strainer. Centrifuge the cell suspension at 350g for 5 min at 4 °C. Discard the supernatant and add 3ml of 1x Red blood lysis buffer for spleen per mouse and incubate at room temperature for 7 min. Add 10 ml of FACS buffer and centrifuge the cells and discard the supernatant. Using CD90.2 microbeads to isolate the CD90.2+ pan T cells. Stain the cells with antibodies of anti-mouse CD4-FITC (11-0042-82, eBioscience) (dilution 1:200) and anti-mouse CD25-PECY7 (557192, BD Bioscience) (dilution 1:200) at 4 °C for 20 mins plus LIVE/DEAD® Fixable Near-IR (L10119, Thermo Fisher Scientific) (dilution 1:500) for dead cell Staining. After staining, sort Tregs (CD4^+^CD25^high^) and conventional CD4^+^CD25^-^ T cells (Tconv) by BD FACSAria™ III sorter. Tconv (1E5) from WT C57BL/6 mice were first labelled with 1 µM CFSE (C34554, Life Technology). Tregs were sorted from *Dj-1*^*-/-*^ mice or wild-type (WT) littermates. Splenocytes depleted of T cells by using CD90.2 beads (130-049-101, Miltenyi Biotec) were irradiated with a total dose of 30 Gy by RS2000 (Rad Source Technologies) and used as feeder cells (2E5). Tconv cells were co-cultured with irradiated feeder cells and Treg cells from various genotypes in the presence of 1 µg/ml soluble anti-CD3 antibody (no azide/low endotoxin, 554829, BD Biosciences). The mice T-cell culture media is the complete RPMI media, which included RPMI 1640 medium (21870084, Thermo Fisher Scientific) supplemented with 10% heat-inactivated fetal bovine serum (FBS, 10500-064, Thermo Fisher Scientific), 50 U/ml penicillin, 50 µg streptomycin (15070-063, Thermo Fisher Scientific), 1 mM sodium pyruvate (11360070, Thermo Fisher Scientific), 2 mM GlutaMAX (35050061,Thermo Fisher Scientific), 0.1 mM non-essential amino acids (M7145, Sigma Aldrich), 50 µM beta-mercaptoethanol (M7522, Sigma Aldrich) and 10 mM HEPES (15630080, Thermo Fisher Scientific). All the mice cells were cultured in 37°C 5% CO2 incubators, unless specified. Gated CD4+ viable single CFSE-labelled cells were measured three days later in a BD LSRFortessa™. The viability of cells was assessed by using the LIVE/DEAD® Fixable Near-IR Dead Cell Stain kit (L10119, Thermo Fisher Scientific) (dilution 1:500).

#### EAE model

The EAE model was performed as previously described ^49^. In brief, each mouse was injected subcutaneously (s.c.) with 115 µg MOG35-55 peptide (Washington Biotech) emulsified in CFA (263810, Difco) plus an intraperitoneal (i.p.) injection of 300 ng pertussis toxin (NC9675592, List Biological) on days 0 and 2. Clinical signs were assessed daily as described ^83^.

#### Intracellular cytokines measurement

For intracellular cytokine measurement, cells (2E5) from spleen and draining lymph nodes were re-stimulated by 50 ng/ml PMA (Phorbol 12-myristate 13-acetate, P8139, Sigma-Aldrich) and 750 ng/ml ionomycin (I0634, Sigma-Aldrich) in the presence of Golgiplug (555029, BD Biosciences) and Golgistop (554724, BD Biosciences) for 5 hrs in 96-well plates. Following cell surface staining, cells were fixed and permeabilized with Cytofix/Cytoperm buffer (554714, BD Biosciences).

#### qPCR of sorted murine Tregs

Total CD25^high^CD4^+^ Tregs were sorted from spleen of young adult (8-12 wks) and very old (∼75 wks) WT *Dj-1*^+/+^ mice. RNA was extracted by using the RNeasy Mini Spin Kit (74104, Qiagen) including column-genome DNA removal steps. cDNA was synthesized by means of the SuperScript III Reverse Transcriptase Kit (18080-044, Invitrogen). qPCR was performed with the LightCycler 480 SYBR Green I Master Mix (4707516001, Roche Applied Science) and on the LightCycler 480 (384-well) real-time PCR platform (Roche) as previously described ^41^. The housekeeping gene for detecting *Dj-1* mRNA expression (QT00167013, Qiagen) in total Tregs was *Rps13* (QT00170702, Qiagen).

#### Microarray analysis

RNA samples of CD25^high^CD4^+^ T cells (Tregs) sorted from around 45-week-old *Dj-1*^*-/-*^ mice and *Dj-1*^*+/+*^ littermates, were analysed with the Affymetrix mouse Gene 2.0 ST Array at EMBL Genomics core facilities (Heidelberg). Samples were first checked with an Agilent Bioanalyzer 2100 by using the RNA 6000 Pico-kit (50671513, Agilent) and only RNA samples with RIN higher than 8.5 were further processed for microarray measurement. The expression signal at the exon level was summarized by the Affymetrix PLIER algorithm ^84^ with DABG and PM-GCBG options by means of the sketch-quantile normalization approach (Affymetrix Expression Console v1.4). The corresponding probesets were considered differentially expressed if they passed the following combinatory filters ^85^: (a) whether the change folds were >= 2 between the means of DJ-1 KO and WT Tregs or Teffs; (b) whether the P-value, resulting from a two-tailed Student t-test, was <=0.05; (c) whether the cross-hyb type of the probeset was equal to 1; (d) whether the probeset with the highest expression level was higher than 100 (with the median value of ∼90 for each our sample); (e) for a given group (e.g. WT) with the higher mean intensity value of the probeset, whether the probeset in all the replicates of the given group was detected as ‘present’ according to the default setting of the Affymetrix Expression Console. DAVID v6.7 was used to perform functional enrichment analysis for the lists of differentially up- or down-regulated genes ^86^. To simplify the analysis of the DAVID enrichment results, we mainly selected representative GO-terms and KEGG pathways with a P-value <0.05 and fold enrichment >=1.5 from each enrichment cluster. Heatmaps were generated using the R Heatmap.2 (gplots package) with scaled row values (with mean 0 and standard deviation 1). The microarray data were deposited into the GEO series database (<u>GSE115269</u>).

#### Seahorse metabolic assay

CD4^+^CD25^high^ Tregs and CD4^+^CD25^-^ Tconv cells from the spleen of old *Dj-1* (∼45 wks) KO and WT mice were sorted on BD FACSAria™ III. 4E5 freshly isolated cells were allowed to rest for 3 hrs in the complete RPMI media before being plated in wells by using Cell-TAK (354240, Corning). Tregs were pooled together from different mice of the same group to reach 4E5/well due to a limited number of Tregs in each individual mouse. The oxygen consumption rate (OCR) was measured in XF base medium (102353-100, Agilent Technologies, from the Seahorse XF Cell Mito Stress Test Kit), containing 1 mM pyruvate (S8636, Sigma-Aldrich), 2 mM L-glutamine (G8540, Sigma-Aldrich) and 25 mM glucose (G8769, Sigma-Aldrich), under basal conditions and in response to 1 µM Oligomycin (103015-100, Agilent Technologies), 1.5 µM FCCP (carbonyl cyanide-4 (trifluoromethoxy) phenylhydrazone) (103015-100, Agilent Technologies) and 1 µM Rotenone and 1 µM Antimycin A (103015-100, Agilent Technologies) by the Seahorse XF96 analyser (Agilent). The results were analysed by Wave 2.6.0 (Agilent Technologies).

#### Treg cell proliferation and iTreg differentiation assay

CD4 T cells were first enriched by anti-mice CD4 (L3T4) microbeads (130-117-043, Miltenyi Biotec) from spleen and peripheral LNs following manufacturer’s instructions. Stain the CD4 T cells with antibodies of anti-mouse CD4-FITC (11-0042-82, eBioscience) (dilution 1:200), anti-mouse CD25-PECY7 (557192, BD Bioscience) (dilution 1:200), anti-mouse CD44-PE (560569, BD Bioscience) (dilution 1:200) and anti-CD62L-PerCP-CY5.5 (560513, BD Bioscience) (dilution 1:200) plus LIVE/DEAD® Fixable Near-IR (L10119, Thermo Fisher Scientific) (dilution 1:500) for dead cell Staining. After staining, sort the naïve CD4 T cells (CD4^+^CD25^-^CD62L^high^CD44^low^) on BD FACSAria™ III sorter.

Naïve CD4 T cells were isolated from spleen and peripheral LNs of *Dj-1*^*+/+*^ WT B6 mice aged 8-12 weeks old. Add 1 µl of 5 mM CellTrace™ Violet dye (CTV, C34557, Thermo Fisher Scientific) into 1 ml of PBS and use it to suspend naïve CD4 T cells. Incubate the cell mixture at 37°C for 7 min and then wash the cells with complete medium for three times. 1.5E5 naïve CD4 T cells were seeded in one well of 48-w plate with flat bottom coated 2 μg/ml of anti-mouse CD3 (553057, BD Bioscience) and 2 μg/ml of anti-mouse CD28 (553294, BD Bioscience) antibodies. Add 2ng/ml human TGF-β (100-21, Peprotech) and 10U/ml mouse IL-2 (212-12, Peprotech) for iTreg differentiation. Cells were treated with PDH inhibitor (CPI-613) (SML0404, Sigma Aldrich) with different concentrations or the vehicle (DMSO, D2650, Sigma Aldrich) or dichloroacetate sodium (DCA, 347795, Sigma Aldrich). Cells were incubated in the incubator for 3 days. On day 2, exchange half of the medium with cytokines and the inhibitor CPI-613 or DCA. On day3, harvest the cells and perform the staining for FACS analysis.

For nTregs proliferation, CD4^+^CD25^high^ nTregs were isolated from the spleen and peripheral LNs of WT B6 mice aged 8-12 weeks old and labelled by CellTrace™ Violet dye. 1E5 nTregs were seeded into 96-well round bottom plate coated with 1 μg/ml of anti-CD3 and 1 μg/ml of anti-CD28 antibodies. 50 U/ml of IL2 was added for nTreg proliferation. On day3, harvest the cells and perform the staining for FACS analysis.

#### Statistical analysis

P values were calculated with non-paired two-tailed Student t test (Graphpad prism or Excel) as specified in Figure legend. The EAE clinical scores between WT and KO groups were compared by paired two-tailed Student t test (Graphpad). All error bars represent the standard deviation. The P-values associated with Pearson correlation analysis were from a two-tailed test generated by GraphPad Prism.

#### Data availability

The microarray data used in this study have been deposited in the Gene Expression Omnibus (GEO) under the access number of <u>GSE115269</u>. To review, go to https://www.ncbi.nlm.nih.gov/geo/query/acc.cgi?acc=GSE115269 Enter the token **qlqbmoqmpryvxkp** into the box.

The proteomic mass spectrometry data [<u>doi:10.25345/C5437F</u>] of the Co-IP analysis has been deposited in ProteomeXchange via MassIVE ^87^ under the identifier PXD016610. To review, go to https://massive.ucsd.edu/ProteoSAFe/static/massive.jsp (login in the top right of the webpage) User: **MSV000084659_reviewer**; Password: **ISB.FengHe.2019**

## Expanded view figures and tables

Expanded view (EV) figures and tables are attached in the end of the manuscript and will be available online.

## Author contributions

E.D. and N.Z. designed and performed major parts of the healthy human- and mouse-related experiments, respectively. N.Z. analyzed mouse-related data. C.C. performed healthy human-, patient- and mouse-related experiments. N.P. and C.L.L. performed and analyzed metabolome-related experiments. D.G.F. and S.D. performed parts of primary human T-cell related experiments. G.G.G. and J.C.S contributed to the seahorse assays. M.G. and J.R. performed and analyzed proteomic experiments. H.K., M.G. and D.B. performed EAE-related experiments. C.G. supervised the flow-cytometry related experiments. D.C. and C.L. supervised mouse-related experiments. D.M.V.W. performed part of EAE experiments. W.W., R.K., R.B. and M.O. provided insights and supervised parts of the experiments. R.B. and M.O. revised the manuscript. F.H. conceived and directed the project and wrote the manuscript.

## Acknowledgements

This work was supported by the Luxembourg National Research Fund (FNR) CORE program grant (CORE/14/BM/8231540/GeDES), Luxembourg-RIKEN bilateral program ‘TregBar’, ‘NEXTIMMUNE’ and ‘CRITICS’ DTU to F.H., individual Aide à la Formation Recherche (AFR) grants to E.D. (PHD-2014-1/7603621) and N.Z. (PHD-2015-1/9989160) and other PhD position grants to C.C. (through PRIDE/2015/10907093) and D.F. (through Luxembourg-RIKEN bilateral/2015/11228353, ‘TregBar’), by the DFG (Grant WU 164/5-1) to W.W., and by the German Federal Ministry of Education and Research (BMBF) through the Integrated Network MitoPD (grant 031A430E) to W. W. and D.M.V.W. We thank Annegrät Daujeumont, Nassima Ouzren, Ting Li, Olga Boyd, Nicolas Bonjean, Caroline Davril, Sandra Köglsberger, Jonas Walter, Susanne Badeke and Fred Fack for their expert technical support. We acknowledge Marie Paule Dufresne (Liege, Belgium) for allowing us to access the irradiator and Nicolas Malvaus from Luxembourg Red Cross for providing buffy coats and leukopaks.

## Competing financial interests

The authors declare no competing financial interests.

## Expanded view (EV) figures

**Figure EV1.**
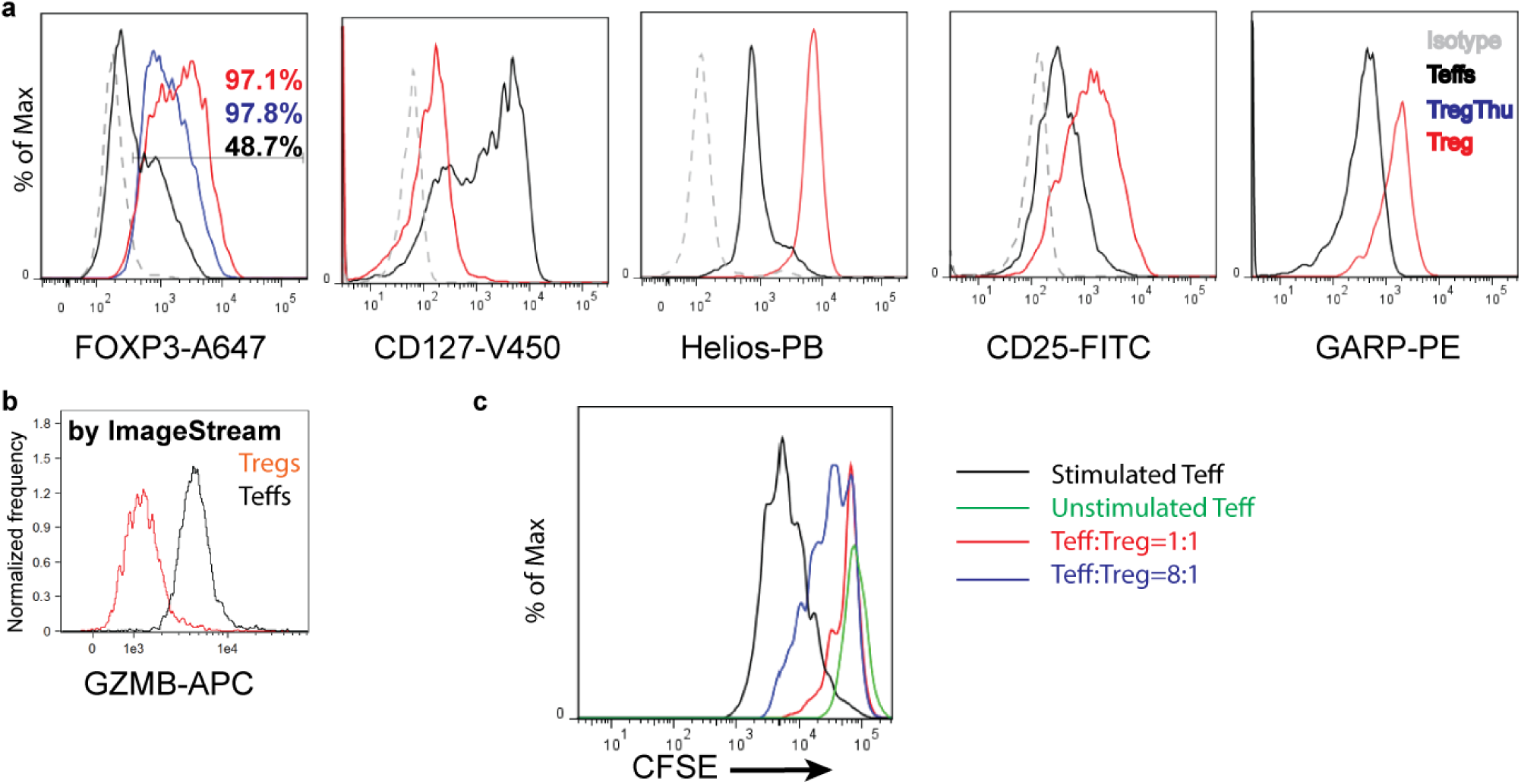
Charaterization of highly purfied human nTregs. **a**, Characterization of highly purified human nTregs (CD4^+^CD25^high^CD127^low^). Representative flow-cytometry plots of FOXP3, CD127, Helios, and CD25 on human Tregs relative Teffs (the markers were not stained in the panel). ‘TregThu’ is a golden standard of isolated human Tregs (refer to Methods and the previous work ^80^). The expression of *LRRC32 (*GARP) was measured after 3-day stimulation by irradiated EBV-transformed B cells. **b**, Representative ImageStream plot of GZMB expression in unstimulated Tregs or Teffs ^88^. **c**, Representative *In-vitro* proliferation assay of human Teffs suppressed by human Tregs at different ratios, in coculture with irradiated EBV-B cells for 5 days. Results represent 7-10 (**a-c**) independent experiments (from different heathy donors).

**Figure EV2.**
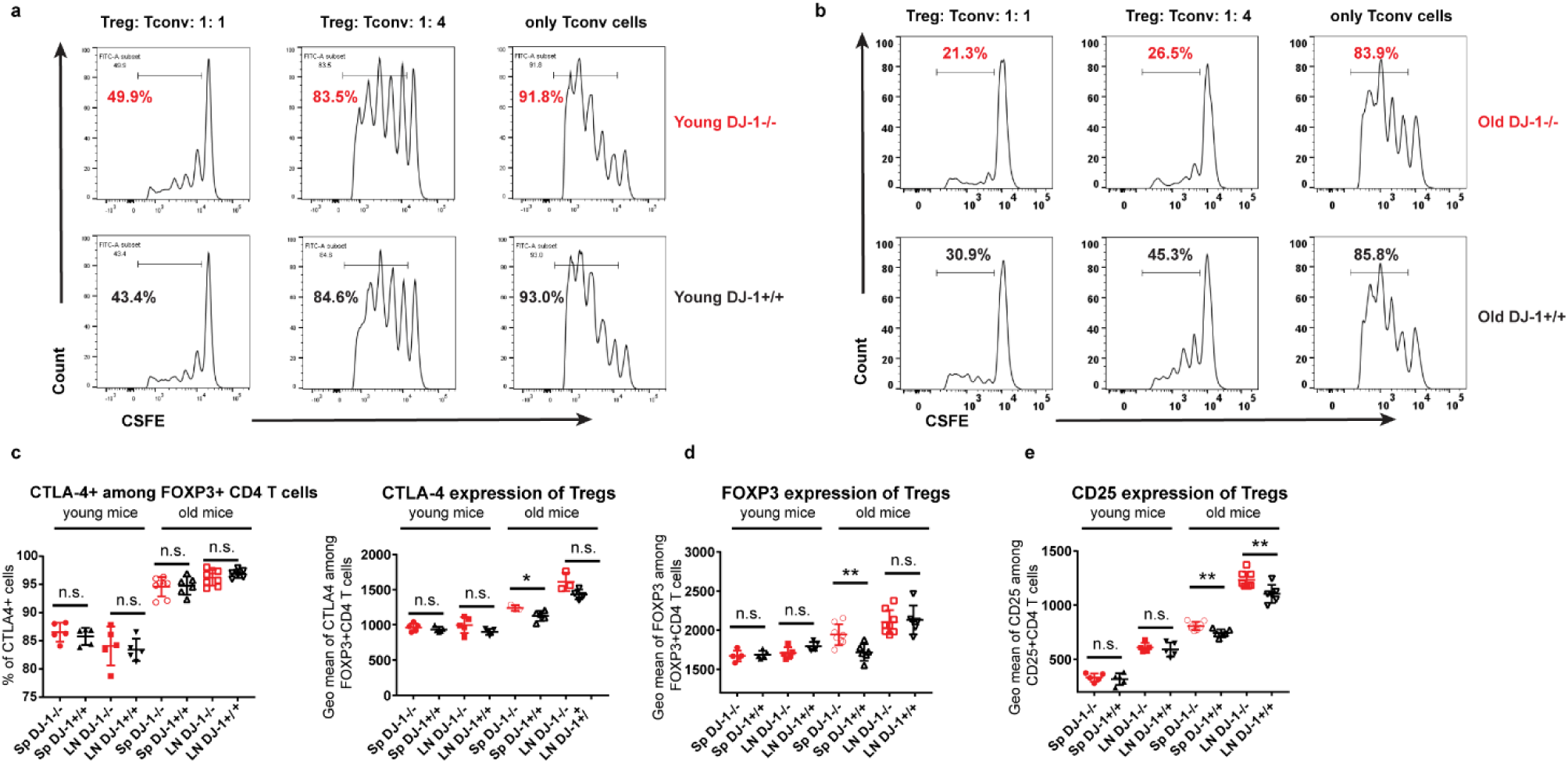
*Dj-1* depletion did not compromise individual Treg suppressor function (TSF). **a, b**, *In-vitro* suppression assay of *Dj-1*^*-/-*^ or *Dj-1*^*+/+*^ Tregs isolated from young (**a**) or old (**b**) mice in coculture with Teff cells and irradiated feeder cells in the presence of anti-CD3 for 3.5 days. Percentage of dividing cells from the total population (the total population indicates the gated DAPI-negative and CD4^+^, CFSE stained Ths) is presented on the plots. **c**, Frequency of CTLA4-expressing cells among CD4^+^FOXP3^+^ Treg cells (left) and Geometric mean of CTLA4 among CD4+FOXP3+ Tregs from spleen (Sp) and lymph nodes (LN) from *Dj-1*^*-/-*^ and *Dj-1*^*+/+*^ WT littermates. **d, e**, Geometric mean of FOXP3 and CD25 expression among CD4+FOXP3+ Tregs. Young/old mouse results are marked in each panel (young KO, n=5; young WT, n=5; old KO, n=8; old WT, n=6; for results of old mice except for CTLA4 Geomean measurement (old KO, n=3; old WT, n=5 sp; old WT, n=4 LN; old KO, n=3 LN; of note, for comparability, the CTLA4 Geomean results were not pooled from different batches), data pooled from 2 independent experiments). Results represent three (**a, b**) and four (**c, d, e**) independent experiments. Data are mean ± s.d. The P-values are determined by a two-tailed Student’s *t*-test. ns, not significant, *P<=0.05, **P<=0.01 and ***P<=0.001.

**Figure EV3.**
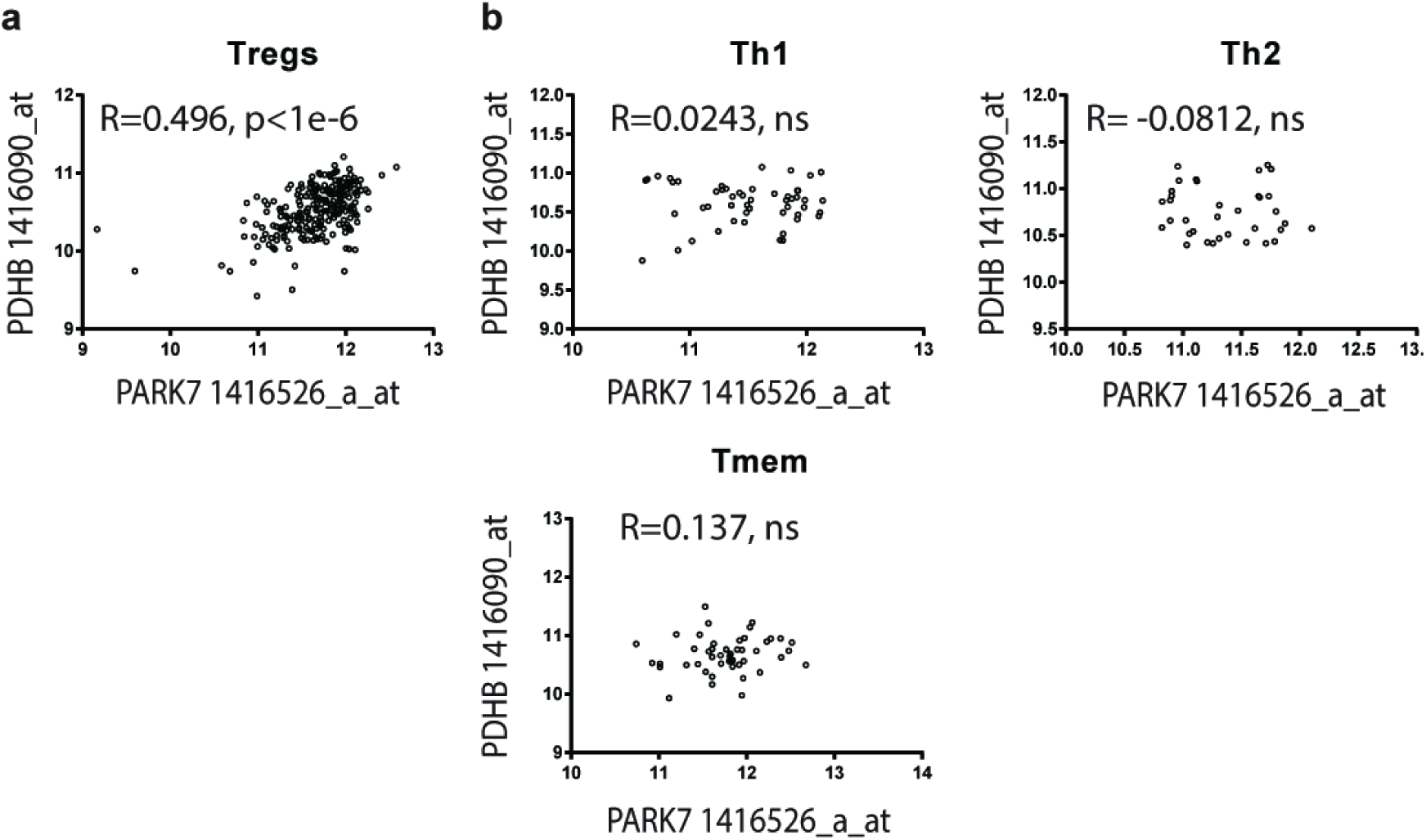
Treg-specific correlation between *Dj-1* and *Pdhb*. **a**, mRNA expression of *Park7*/*Dj-1* is highly correlated with *Pdhb* only in murine Tregs but not in other tested CD4 T cell subsets. The number of microarray samples used for correlation analysis is 240, 51, 35 and 47 from Tregs, Th1, Th2 and Tmem (memory CD4 T cells) respectively. The R value corresponds to Pearson correlation coefficient and the associated P-value is from a two-tailed test (Prism). Ns, non-significant and otherwise, the P-value is provided for the corresponding cell type. For the details, refer to the database (from *Sakaguchi’s lab*): https://sysimm.ifrec.osaka-u.ac.jp/immuno-navigator/?o=10090&probe1=1416526_a_at&probe2=1416090_at&cell=5.

**Figure EV4.**
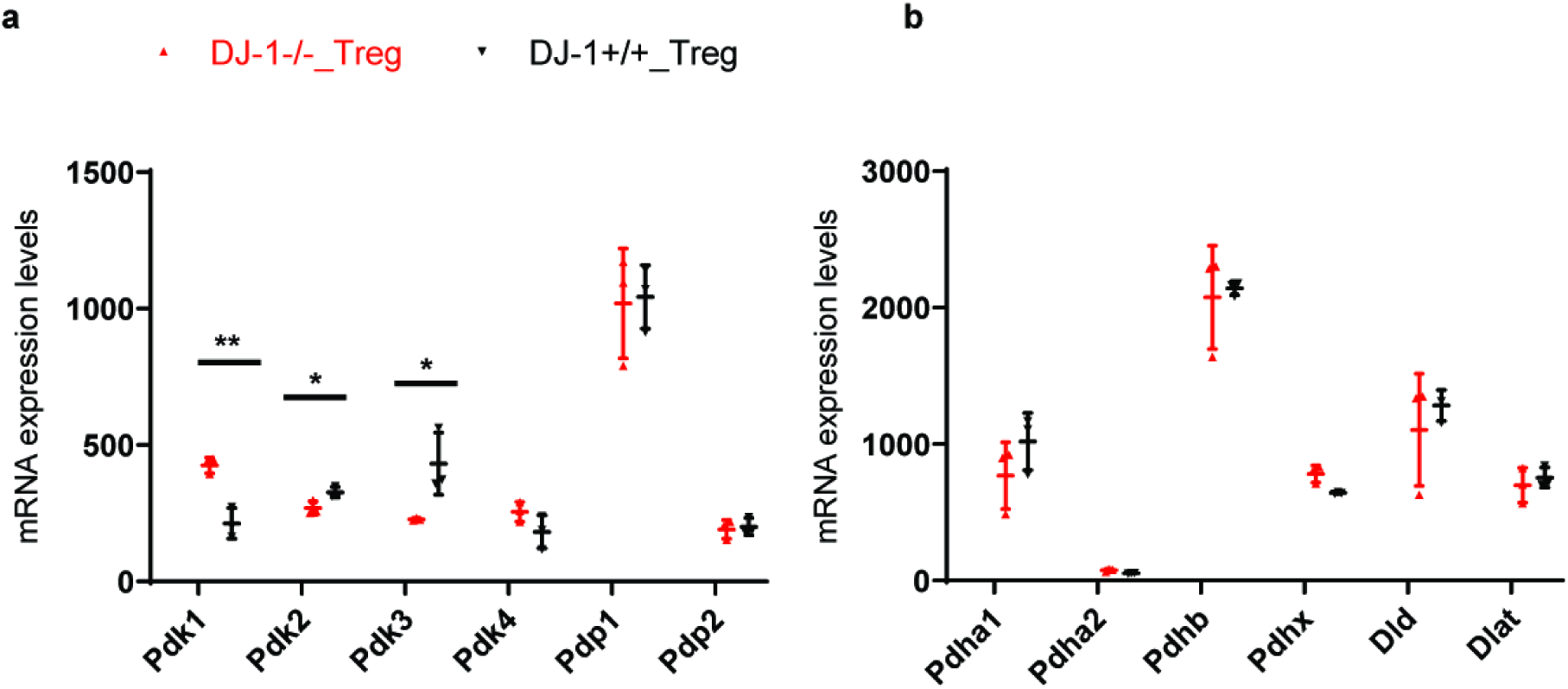
Expression analysis of PDH complex encoding and regulatory machinery genes in Tregs of old mice. **a**, mRNA expression of PDH kinases (*Pdks*) and pyruvate dehydrogenase phosphatases (*Pdps*) in Tregs freshly isolated from old Dj-1 KO and WT littermates. **b**, mRNA expression of several PDH complex encoding genes in Tregs freshly isolated from old *Dj-1 KO* and WT littermates. n=3 per group of mice. Results are the summary of the microarray datasets. Data are mean± s.d. The P-values are determined by a two-tailed Student’s *t*-test. ns or unlabelled, not significant, *P<=0.05, **P<=0.01 and ***P<=0.001.

## Expanded View (EV) Tables

**Table EV1.**
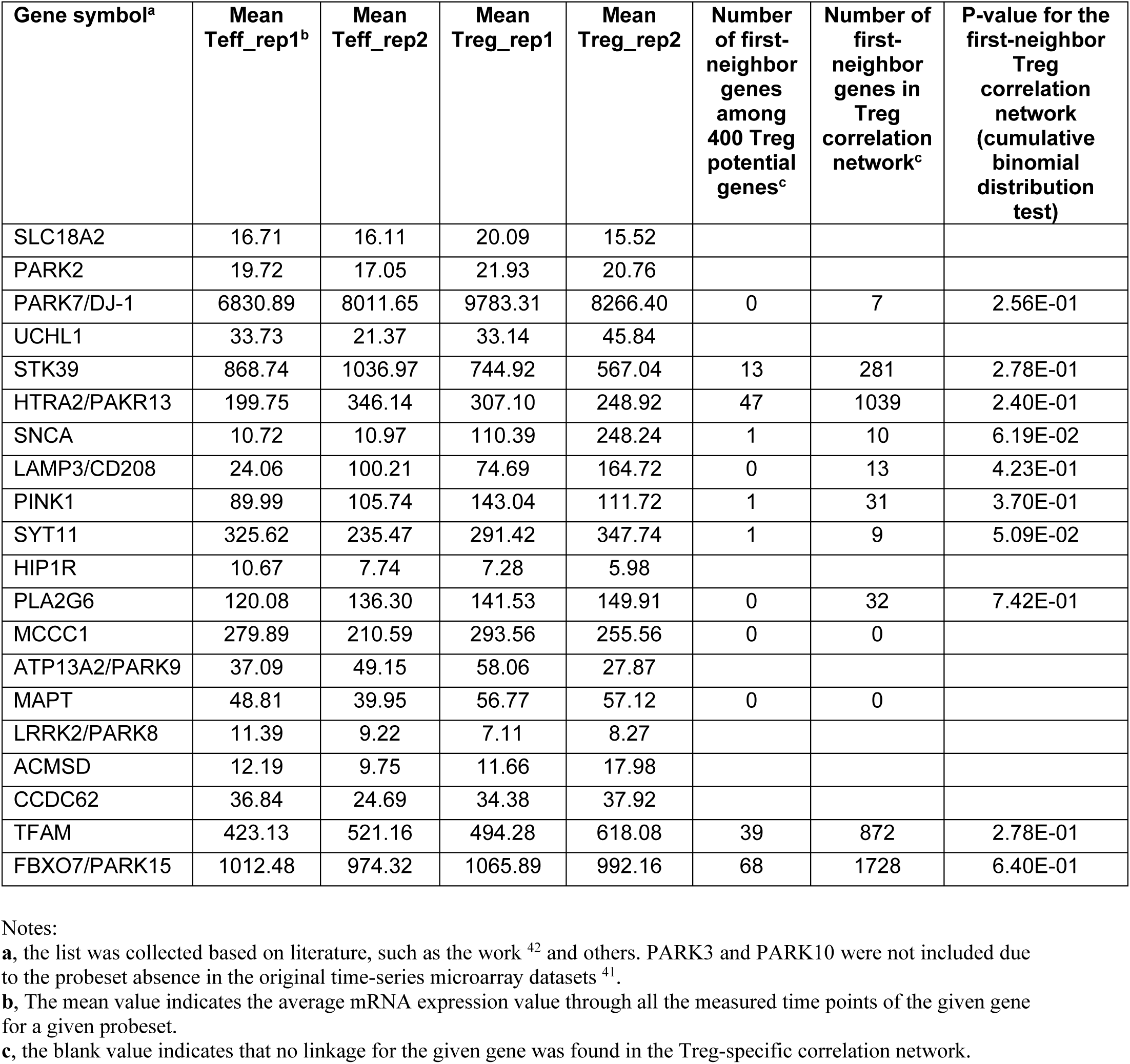
The known PD-related genes measured in published human Treg time-series microarray datasets ^41^.

**Table EV2.** The Co-IP-MS identified potential binding partners of DJ-1 in human primary Tregs/Teffs.

| Gene symbol | Mean<br>Teff_rep1 | Mean<br>Teff_rep2 | Mean<br>Treg_rep1 | Mean<br>Treg_rep2 | Fold change<br>(Tregs sti vs.<br>unsti)<sup>b</sup> | Fold change<br>(Tregs sti vs<br>unsti (without<br>IL-2)) | Fold change<br>(Teffs sti vs.<br>unsti) | Fold change<br>(DJ-1 ab vs<br>IgG in Teffs) | Binding<br>preference<sup>c</sup> |
| --- | --- | --- | --- | --- | --- | --- | --- | --- | --- |
| PDHB | 1304<br>.83 | 1248.<br>61 | 1650.1<br>3 | 1331.09 | 198.13 | 2.43 | 70.53 | 64.01 | unsti |
| GNB2L1 | 8549<br>.45 | 14203<br>.38 | 11974.<br>55 | 9578.58 | 36.47 | 295.49 | 16.63 | 35.09 | sti |
| EXOSC1 | 425.<br>11 | 452.6<br>9 | 736.40 | 555.02 | 5.11 | 2.17 | 66.07 | 33.03 | sti |
| EIF2A | 3478<br>.47 | 3370.<br>57 | 2809.8<br>5 | 2844.83 | 3.33 | 22.16 | 9.86 | 25.07 | sti |
| SART1 | 253.<br>46 | 335.8<br>2 | 279.24 | 265.06 | 13.87 | 63.89 | 29.02 | 19.17 | sti |
| ELAVL1 | 788.<br>85 | 1908.<br>74 | 1253.6<br>9 | 999.38 | 19.02 | 117.64 | 46.47 | 17.40 | sti |
| ATP1A1 | 489.<br>10 | 1548.<br>97 | 891.64 | 615.27 | 7.43 | 21.02 | 4.06 | 14.48 | unsti |
| SKIV2L2 | 639.<br>83 | 703.7<br>1 | 717.41 | 794.02 | 28.65 | 34.13 | 89.77 | 9.86 | sti |
| DOCK10 | 1883<br>.18 | 1345.<br>03 | 1584.8<br>8 | 1580.57 | 2.11 | 3.79 | 2.89 | 8.51 | sti |
| ATP5D | 273.<br>57 | 687.9<br>8 | 469.23 | 330.71 | 76.51 | 99.86 | 3.05 | 7.85 | sti |
| ANXA4 | 359.<br>40 | 660.3<br>6 | 1084.7<br>9 | 955.29 | 7.58 | 14.57 | 2.09 | 6.64 | sti |
| LARS | 597.<br>75 | 744.0<br>1 | 857.39 | 705.88 | 3.88 | 6.06 | 63.76 | 6.49 | unsti |
| HNRNPH<br>3/HNRPH<br>3 | 2211<br>.08 | 1599.<br>24 | 1270.3<br>1 | 1961.07 | 4.63 | 20.33 | 2.61 | 5.26 | sti |
| STT3B | 4887<br>.15 | 4743.<br>66 | 4601.9<br>4 | 5918.17 | 6.32 | 5.45 | 6.73 | 5.18 | unsti |
| EIF5B | 2652<br>.07 | 2294.<br>71 | 1998.9<br>2 | 3190.99 | 5.80 | 3.11 | 11.02 | 5.00 | sti |
| DHX36 | 2543<br>.54 | 1342.<br>67 | 1245.6<br>6 | 1596.94 | 2.26 | 14.27 | 4.74 | 4.95 | unsti |
| KHSRP | 294.<br>26 | 557.8<br>5 | 439.18 | 506.72 | 4.22 | 17.58 | 17.45 | 4.88 | sti |
| ARPC2 | 1139<br>0.36 | 10741<br>.65 | 10140.<br>17 | 8212.70 | 5.37 | 82.96 | 17.28 | 4.88 | unsti |
| PRPF8 | 765.<br>71 | 895.5<br>1 | 784.17 | 603.34 | 4.65 | 9.36 | 11.26 | 3.73 | sti |
| PPP1CC;<br>PPP1CB<sup>a</sup> | 6423<br>.858 | 5781.<br>311 | 6891.0<br>76 | 8885.82<br>1 | 2.80669<br>1 | 2.65 | 17.76 | 3.16 | sti |

Notes:
a, the mean mRNA expression value in Tregs/Teffs was taken from PPP1CB. The mean expression intensity values through the first 6hr period following the TCR stimulation were taken from the previous time-series microarray dataset <sup>41</sup>. b, the short term “unsti” and “sti” represents the word “unstimulated” and “stimulated”, respectively c, for those bound preferentially in unstimulated T cells, the reciprocal fold changes are shown (i.e., 1/x).

**Table EV3.** List of confirmed known DJ-1-binding partners (the list of known DJ-1 binding partners were downloaded from NCBI or pubmed literatures).

| Gene symbol | Mean<br>Teff_rep1 | Mean<br>Teff_rep2 | Mean<br>Treg_rep1 | Mean<br>Treg_rep2 | Fold change<br>(Treg sti vs.<br>unsti) | Fold change<br>(Treg sti vs<br>unsti (without<br>IL2)) | Fold change<br>(Teff sti vs.<br>unsti) | Fold change<br>(Jurkat cells sti<br>vs. unsti) | Fold change<br>(DJ-1 ab vs IgG<br>in unsti Teff) | Fold change<br>(DJ-1 ab vs IgG<br>in unsti Jurkat) |
| --- | --- | --- | --- | --- | --- | --- | --- | --- | --- | --- |
| HDAC1 | 981.49 | 2009.7<br>7 | 2142.04 | 1322.4<br>5 | 192.53 | 1.11 | 1.36 | 0.78 | 2.34 | 12.02 |
| NPM1 | 4093.4<br>9 | 4232.0<br>0 | 5464.82 | 5882.7<br>1 | 35.58 | 1.55 | 0.16 | 1.60 | 17.62 | 131.70 |
| PARK7 | 6830.8<br>9 | 8011.6<br>5 | 9783.31 | 8266.4<br>0 | 0.71 | 0.95 | 2.25 | 1.18 | 2.66 | 56.98 |
| TALDO1 | 1291.2<br>6 | 2576.7<br>5 | 2603.60 | 1181.2<br>3 | 0.04 | 2.70 | 2.70 | 0.06 | 4.56 | 3.54 |
| PRDX2 | 1220.9<br>1 | 3986.8<br>4 | 2964.41 | 1512.8<br>4 | 0.01 | 0.13 | 1.40 | 0.98 | 1.78 | 3.69 |
| CHD4 | 780.61 | 622.41 | 673.84 | 560.56 | 28.62 | 49.22 | 1.37 | 0.83 | 13.32 | 31.95 |
| RBBP4 | 893.34 | 1434.2<br>2 | 838.35 | 788.77 | 0.54 | 1.05 | 2.44 | 0.92 | 4.22 | 21.06 |
| MTA1 | 334.51 | 518.85 | 388.81 | 324.99 | 3.32 | 39.47 | 0.72 | 0.76 | 38.24 | 165.48 |
| GATAD2<br>B | 256.19 | 183.64 | 203.31 | 257.81 | 0.62 | 1.13 | 0.95 | 0.75 | 10.00 | 26.41 |
| MTA2 | 8.15 | 7.00 | 9.91 | 4.45 | 1.59 | 0.92 | 1.42 | 0.82 | 4.92 | 28.43 |
| STAT1 | 2473.8<br>79 | 3330.8<br>37 | 5127.19<br>2 | 6578.4<br>21 | 10.57 | 0.053 | 0.86 | 1.49 | 42.29 | Binding<br>to igG in<br>Jurkat |

Notes: The list included the genes independent of their mean mRNA expression levels. Here only the reported binding partners of DJ-1 confirmed by our MS analysis were included. The Jurkat cells were stimulated by PMA (10 ng/ml) and ionomycin (1 µg/ml) for 4 h. The human primary T cells were stimulated as described in the Materials and Methods.

